# A tradeoff between enterovirus A71 particle stability and cell entry

**DOI:** 10.1101/2022.09.28.506941

**Authors:** Adam Catching, Ming Te Yeh, Simone Bianco, Sara Capponi, Raul Andino

## Abstract

A central role of viral capsids is to protect the viral genome from the harsh extracellular environment while facilitating initiation of infection when the virus encounters a target cell. Viruses are thought to have evolved an optimal equilibrium between particle stability and efficiency of cell entry. In this study, we genetically perturbed this equilibrium in a non-enveloped virus, enterovirus A71 to determine its structural basis. We isolated a single-point mutation variant with increased particle thermotolerance and decreased efficiency of cell entry. Using cryo-electron microscopy and molecular dynamics simulations, we determined that the thermostable native particles have acquired an expanded conformation that results in a significant increase in protein dynamics. Examining the uncoating intermediate states of the thermostable variant suggests a pathway, where the lipid pocket factor is released first, followed by internal VP4 and finally the viral RNA.

## Introduction

All proteins must form one or a continuum of native, folded, states to perform their required functions, and they are often required to traverse numerous semi-folded metastable states before reaching the low-energy “ground-state”^1^. Molecular chaperones and specific cellular conditions may aid in the proper folding pathway^2,3^, and chemical, physical, and cellular environments also catalyze the reversion to partially unfolded or even fully denatured states. Proteins also have dynamic ranges in which they are natively folded and/or functional and that may include large temperature ranges^4^.

Proteins of virus particles constitute an interesting special case: they must resist harsh extracellular conditions, while evading detection and degradation within the infected organism^5,6^. Accordingly, the central role of virus particles is to protect the viral genome during virus spread within and between individuals. Most importantly, the capsid must also allow the effective release the genome at the right time to initiate viral infection. The viral capsid traverses a potentially large landscape of environments and conformational states, from assembly within the cell^7,8,9^ to the highly variable extracellular environment to the cell entry pathway^10,11^.

Viruses have evolved efficient cell entry mechanisms^12,13,14^ that depend on effective genome release triggered by interactions with specific virus receptors, often facilitated by optimal pH within the endosomal compartment, and membrane proximity^15,16,17^. Within a simpler experimental paradigm, genome release can be simulated by heating purified virus particles^18,19,20^. Enteroviruses particles are composed of single-strand RNA (virion RNA) genomes encapsulated by an icosahedral capsid of approximately 30 nm in diameter^21^. While the 20^th^ century was marked by the near eradication of poliovirus, a prototypical Enterovirus, outbreaks of non-poliovirus enteroviruses have become a topic of concern^13,22^. As one recent example, Enterovirus A71 (EV-A71) is a global infectious disease that in recent years has affected the Asia-Pacific region causing several outbreaks^23^. The virus is the main etiological agent for hand, foot, and mouth disease with global outbreaks and epidemics. Infection causes severe neurological, cardiac, and respiratory problems in young children.

In general, the enormous adaptation capacity of RNA viruses, such as EV-A71, is powered by their ability to acquire new mutations under the proper conditions or challenges. RNA-dependent RNA polymerases of RNA viruses exhibit characteristically low fidelity with measured mutation rates of 10^-3^ to 10^-5^ mutations per nucleotide copied per replication cycle^24,25^. These mutation rates are orders of magnitude greater than those of nearly all DNA-based viruses and organisms. Given the short viral generation times and large population sizes in experimental and natural infections, every possible point mutation and many double mutations may be generated with each viral replication cycle and may be present within the population at any time^26^. Because RNA viruses exist as swarms of similar variants that are continuously regenerated by mutation of related sequences, strong selection pressures result in steeper fitness landscapes and influence which mutations become fixed in the population^27^.

In this study, we exploited the great adaptive capacity of EV-A71 to isolate variants with increased particle thermostability. This experimental strategy enabled us to examine biophysical and structural flexibility of enterovirus particles and to better understand the specific steps that the virus undergoes during the entry process. Surprisingly, even if the isolated variant is less efficient to enter cells, in an animal model of infection, the thermotolerant virus is more neurovirulent, indicating that capsid conformation plays an important role in tissue tropism and pathogenesis.

## Results

### Identification of EV-A71 thermostable variants

To examine the relationship between particle stability and infectivity and to obtain empirical support to this prediction, we conducted a genetic experiment to isolate thermostable-particle variants. First, we studied the effect of temperature in particle inactivation. Briefly, we incubated EV-A71 virus stocks (4643/TW/98 strain ~10^5^ TCID_50_) for 1 hour at different temperatures and found that incubation at 46°C reduces titers by 10-fold (Supplementary Fig. 1a). We then carried out a genetic experiment starting from a single laboratory-derived clone EV-A71 4643 (WT). Populations were obtained after serial passages. Eight serial passages were carried out transferring for a single replication cycle (8 hours) in human rhabdomyosarcoma (RD) cell culture (Fig. 1a). To control the influence of drift due to genetic bottlenecks and recombination and complementation between viral variants^28^, each passage was conducted at low multiplicity of infection (MOI= 0.01), using 10^6^ tissue-culture infectious dose 50 (TCID_50_) from the previous passage. Between each passage, we either incubated for 1 hour the supernatants of infected cells at 46°C (passages 1 to 4) and at 48°C (passages 5-8) (Heated Passages) or incubated at 4°C (Control Passages). We then determined the proportion of viruses resistant to heat (i.e., the thermostability of each population) by examining the effect on virus titer of 1-hour incubation at 46°C or 48°C (Fig. 1b). The fraction of thermal resistant viruses for populations selected at higher temperatures (P5 to P8, red) were unaffected by incubation either at 46° or 48°C when comparing control populations (black) (Fig. 1b). Interestingly, control populations were more sensitive to the temperature after 4 rounds of selection. This observation suggests that adaptation to cell culture replication, inversely correlates with thermo-stability.

**Fig. 1.**
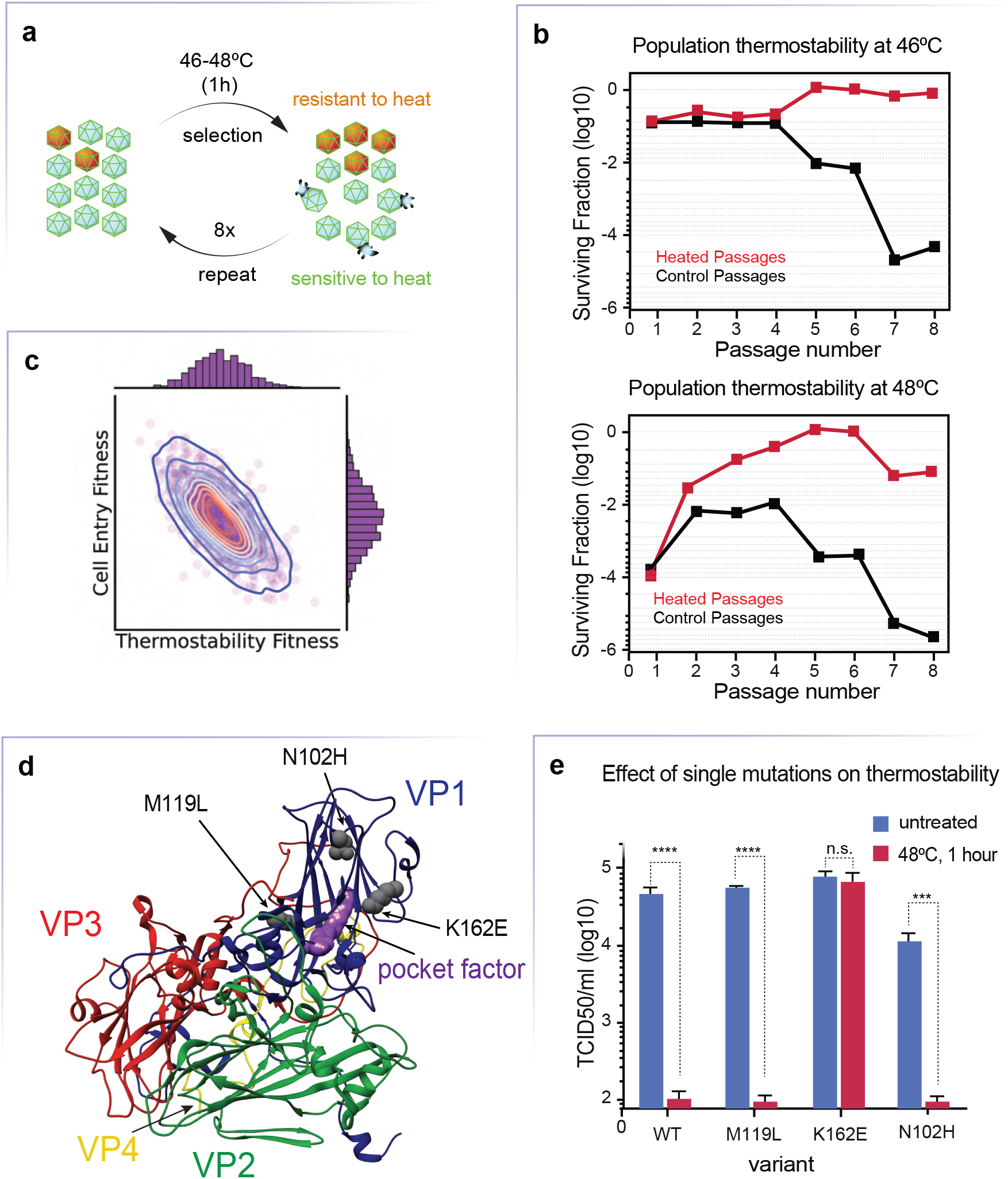
Selecting for thermostable EV-A71 variants. **a**, Computer simulation of virus fitness distributions with different degrees of particle thermostability and cell entry efficiency. The distribution of the virus genotypes determines the fitness relationship between thermostability and infectivity. The simulation predicts that an increase in thermostability corresponds to de-adaptation in the efficiency of virus cell entry. **b**, Experimental design for the selection of virus particles adapted to elevated temperature. **c**, Thermostability assay on virus population selected during serial passage at higher temperature. Each population was evaluated at 46°C (left) 48°C (right). Graphics represent the fraction of infectious viruses resisting 46°C or 48°C treatment (1 hour), relative to the total virus in that population. Red lines are populations selected at a higher temperature; black lines correspond to populations that were not heated between passages. d, Mapping mutations identified in the selected population. Mutations are mapped into the EV-A71 capsomere, subunits VP1 (blue), VP2 (green), VP3 (red), and VP4 (yellow) are shown. **e**, Mutations M119L, N102H and K162E were introduced into the infectious EV-A71 cDNA. Recombinant viruses were assays in cell culture (RB cells) by examining loss of infectivity by treatment at 48°C for 1 hour. ***, p<0.001; ****, p<0.0001.

We hypothesized that adaptation to cell culture replication could result from an increase in the efficiency on uncoating and cell entry. Conversely, adaptation to higher temperature may reduce infectivity. To visually recapitulate this hypothesis, we carried out a numeric experiment assuming that particle stability correlates inversely with uncoating and, thus, virus infectivity^29^. The distribution and dynamics of minor alleles play important roles in population fitness, robustness, adaptation, and disease^30,31,32,33,34^. Virus evolution carried out under relatively simple cell-culture selection pressures follows a quasi-Gaussian distribution of fitness values^35^). This distribution is centered at a relative fitness value of 0.6, plus a significant fraction of deleterious mutations with 0 fitness value, which are considered unviable variants. We simulated the distribution of mutational fitness values by randomly drawing 3000 Normally distributed fitness values centered at 0.6 with a standard deviation of 0.2 for the thermostability phenotype, then calculated the inverse relationship with cell entry (Fig. 1c). Additional random white noise with covariance of 0.05 was introduced to the cell entry fitness values to replicate a non-perfect one-to-one correlation. This procedure allows to visually demonstrate an example of the boundaries of capsid functions controlling stability and uncoating in the context of RNA virus dynamic populations, showing the empirical results indicating that the cell entry fitness distribution is compromised with the increased particle thermal stability (Fig. 1b).

Sequencing of genomic RNA from populations (P6) selected to high temperature identified three mutations mapping within the VP1 coding region: methionine to leucine at position 119 (M119L), lysine to glutamic acid at position 162 (K162E), and asparagine to histidine at position 102 (N102H) (Fig. 1d and Supplementary Fig. 1b). To identify which mutation is responsible for the thermostable phenotype we engineered each mutation into an EV-A71 4643 infectious cDNA. The substitution of basic lysine at position 162 for glutamic acid conferred resistance to 1-hour 48°C incubation (Fig. 1e). In contrast, M119L and N102H did not prevent inactivation at high temperatures. Furthermore, these substitutions were also identified in the control passages where the virus was not subject to high temperatures, suggesting that these mutations are fixed during passages because they confer adaptation to cell culture (Supplementary Fig. 1c). We concluded that a single amino acid substitution, K162E, is responsible for particle thermostability.

### Inactivation is caused by a slowed transition to A-particle

Loss of infectivity by heating results from the capsid protein conformational conversion of a native virus particle to a non-infectious “altered particle” (A-particle)^38^. Conversion to an A-particle in enteroviruses follows a pathway that is thought to represent the conformational intermediates that lead to the physiological virus uncoating and entry process. The A-particle capsid proteins expand outward, the virion RNA genome becomes reorganized, the N-terminus of capsid protein VP1 is exposed on the surface of the particle, and VP4 is lost^11,36,37^.

To examine the biophysical properties of thermostable K162E, we initially employed a fluorescent-based assay, particle stability thermal release assay (PaSTRy)^39^. In this assay, the protein shell reconfigures to render the hydrophobic domains within capsid protein accessible to fluorescent dye. Thus, the interaction of SYPRO orange dye with these exposed hydrophobic regions during thermal conversion of native capsids to A-particles is manifested by an increase in fluorescence. We incubated purified virus particles with SYPRO orange and monitored fluorescence as a function of the temperature increase. The maximal fluorescent peak for wild-type (WT) was around 57.6°C, and for K162E particles the fluorescence peak was shifted 2.4°C (~60 °C) (Fig. 2a). This is consistent with the idea that higher temperature is required for K162E to induce a change in particle conformation. A second signal peak was observed around 75°C, at which temperature additional hydrophobic domains become exposed as capsid proteins undergo additional unfolding.

**Fig. 2.**
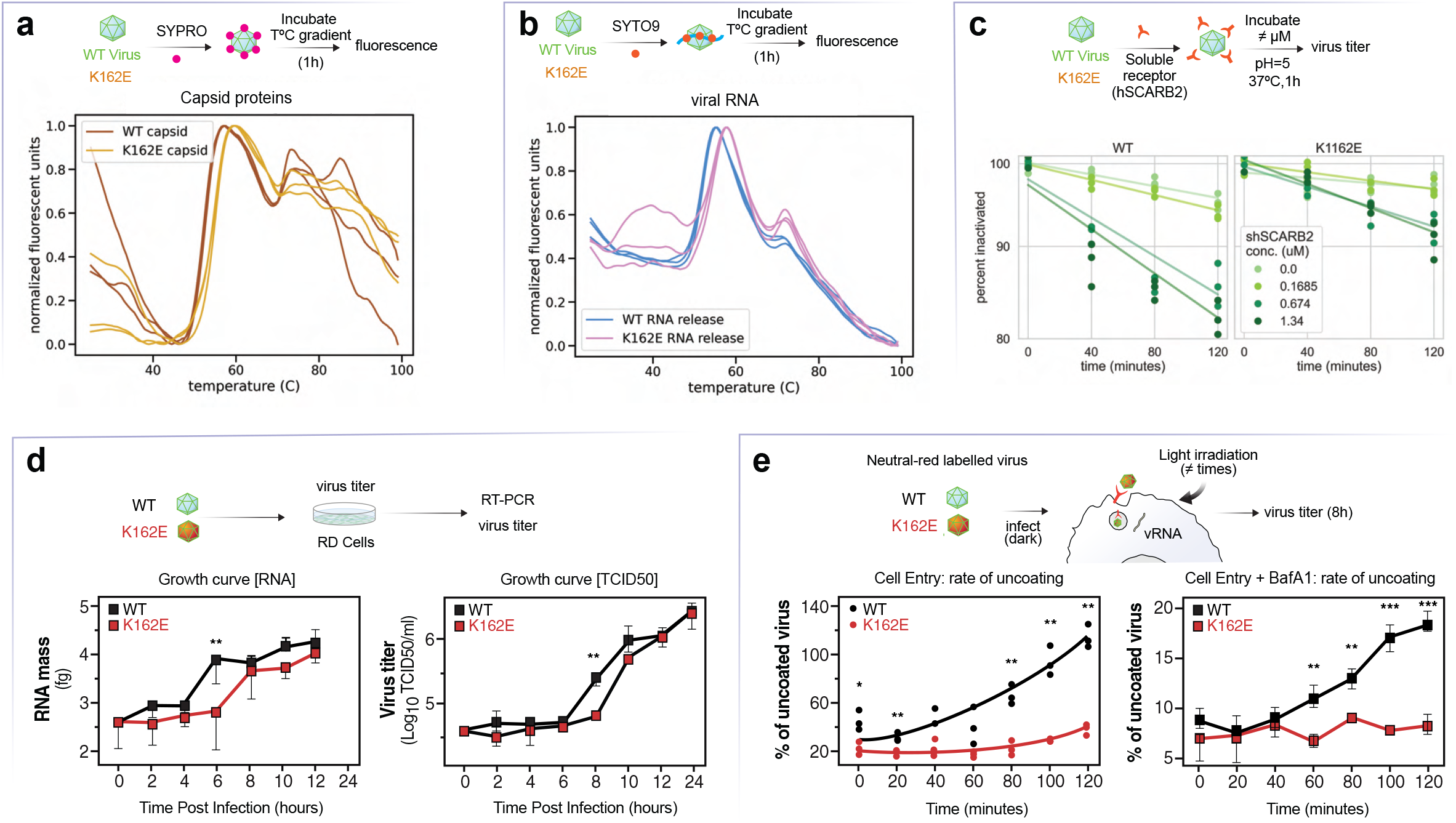
Examining EV-A71 particle conversion. **a** and **b**, WT and K162E variant particle conversion analyzed using fluorescent probes that associate with hydrophobic regions of capsid proteins (SYPRO) or viral RNA (SYTO9) and report on conformational changes in the protein shell or genomic RNA. Changes in fluorescence emission were monitored as a function of the temperature. **c**, WT and K162E variant particle conversion by soluble SCARB2 receptor (shSCARB2) at pH=5. Viruses were incubated for 1 hour at 37°C with increasing concentrations of the shSCARB2, and the proportion of converted particles (virus inactivated) was determined by plaque assay. **d**, One-step growth curve of WT and K162E evaluated by either production of viral genomic RNA or infectious units (TCID_50_). **e**, Left: Virus cellular entry in RD cells determined by the proportion of neutral red-labeled virus that is resistant to illumination at different timepoints after infection (0 to 120 min). Right: BafA1 sensitivity. BafA1 treatment neutralizes pH within the endosome and virus cell entry is delayed. This assay is consistent with the conclusion that K162E demonstrates a defect on cell entry, as it is more sensitive to BafA1 treatment than WT. Significance was calculated using a two-tailed Student’s t-test.*p<0.05, ** p<0.01, *** p<0.001

We further examined particle conversion using a different fluorophore, SYTO9, which monitors release of genomic RNA (Fig. 2b). In this assay, WT releases most of its genome at 51.8°C, and K162E requires 54°C. Thus, these results are consistent with the observation that the K162E variant is more tolerant to temperature than WT (Fig. 1e) and suggest that the transition from native to an A-particle requires additional energy.

During entry, conformational changes leading to uncoating are triggered by cellular factors, such as receptor binding and/or exposure to low pH within endosomes^40^. These conformational changes are thought to induce membrane perturbation necessary for delivery of the viral genome into the cytoplasm of the target cell. The transition of native EV-A71 particle to A-particle can be induced at 37°C by the interaction with a soluble form of the virus receptor, scavenger receptor B2 member 2 (SCARB2), at pH 5^41,42,11^. We compared the rate of the A-particle transition for K162E and WT virus and found that K162E conversion was less efficient than for WT (Fig. 2c). Addition of increasing concentrations of SCARB2 induced inactivation of both WT and K162E, but the proportion of inactivated particles was lower for K162E than WT (Fig. 2c). These results indicate that under temperature and pH physiological conditions, K162E is less likely to undergo the conformational transition required to virus entry.

### Increased capsid stability delays cell entry

Next, we investigated the efficiency of cell entry. First, we compared replication efficiency of the WT and K162E in a one-step growth curve in RD cells. Viral RNA replication was assayed by RT-qPCR every 2 hours (Fig. 2d). WT undergoes an exponential RNA accumulation phase at 4–6 hours post infection (h.p.i.). For K162E, RNA synthesis accumulation was delayed by 2 hours (4–8 h.p.i.). However, similar amounts of viral RNA were detected at 8–12 h.p.i. Virus production was also delayed by 2 hours for K162E, compared to WT. However, at the end of the replication cycle, both viruses had produced similar amounts of viral progeny. These data suggest that cell entry is delayed for K162E, but RNA synthesis and virus production appear to be unaffected.

Neutral red-labeled assay was used to examine the early events of K162E infection. The light-sensitive neutral red–containing EV-A71 was used to distinguish input virus and virus that had initiated replication. The mechanism underlying inactivation is not completely understood. However, once encapsulated into the virion, neutral red is thought to be photoactivated to inactivate the virus via viral genome cross-linking to viral capsid, thus preventing viral uncoating and infection. If the RNA is delivered before light is shined, neutral red is diluted so that it no longer effectively inactivates the virus^43^. Neutral red-labeled WT and K162E were used to infect RD cells, and every 20 minutes, we inactivated a sample by exposing the cells to light, and infection for an already uncoated virus was allowed to proceed for 12 hours. In agreement with our previous results (Fig. 2d), the thermostable mutant initiates infection at a slower rate, with only 40% of K162E virus was able to initiate infection after 2 hours, but 100% WT initiated infection after 100–120 minutes post-infection (Fig. 2e, left panel).

As EV-A71 entry depends on acidification of the endocytic compartment^40^, we next examined virus sensitivity to Bafilomycin A1 (BafA1). BafA1 specifically targets the vacuolar-type H^+^-ATPase (V-ATPase) enzyme, a membrane-spanning proton pump that acidifies either the extra-cellular environment or intracellular organelles, such as the lysosome^44^. To test K162E dependency on low pH, RD cells were pretreated with BafA1 1 hour prior to the addition of neutral red– labeled virus, cells were exposed to light at different times post-infection, and the infection continued for 12 hours at 37°C (Fig. 2e). Under these conditions, WT virus-initiated infection after 60 minutes, and at 120 minutes, 18% of the initial virus had uncoated and initiated infection. In contrast, K162E was unable to infect if the V-ATPase was inhibited by BafA1. Indeed, K162E was approximately 20 time more sensitive to BfaA1 than WT (Supplementary Fig. 2b). These results indicate that K162E is hyper-dependent of lysosome acidification, whereas WT uncoating and initiation of infection can proceed in the presence of BafA1 at a moderate rate.

In addition, the specific infectivity of K162E (e.g., how many viral particles are required per plaque-forming units (pfu)) is 20 time higher than for WT (Supplementary Fig. 2c). These data indicate that the thermo-tolerant K162E variant is less effective at initiating infection.

### Thermodynamic analysis of EV-A71 particle stability reveals the basis for K162E thermo-tolerance

To examine K162E thermodynamic properties of the EV-A71 capsid, we conducted a heat inactivation experiment. Purified K162E and WT were treated for 1 hour at different temperatures, and the reduction of virus infectivity was determined in samples taken every 10 minutes (Fig. 3a). Inactivation followed an exponential decay curve that renders more than 99% of the virus no longer infectious after 1 hour incubation at the highest temperature tested. For WT, incubation at 46° and 48°C inactivated 90% and 99% of the virus, respectively (Fig. 3a). In contrast, K162E was inactivated by incubation at 52°C. While WT survived a maximal incubation temperature of 44°C, K162E did not lose infectivity at 50°C. Thus, a single point mutation at position K162 of VP1 increases temperature resistance by 4–6°C.

**Fig. 3.**
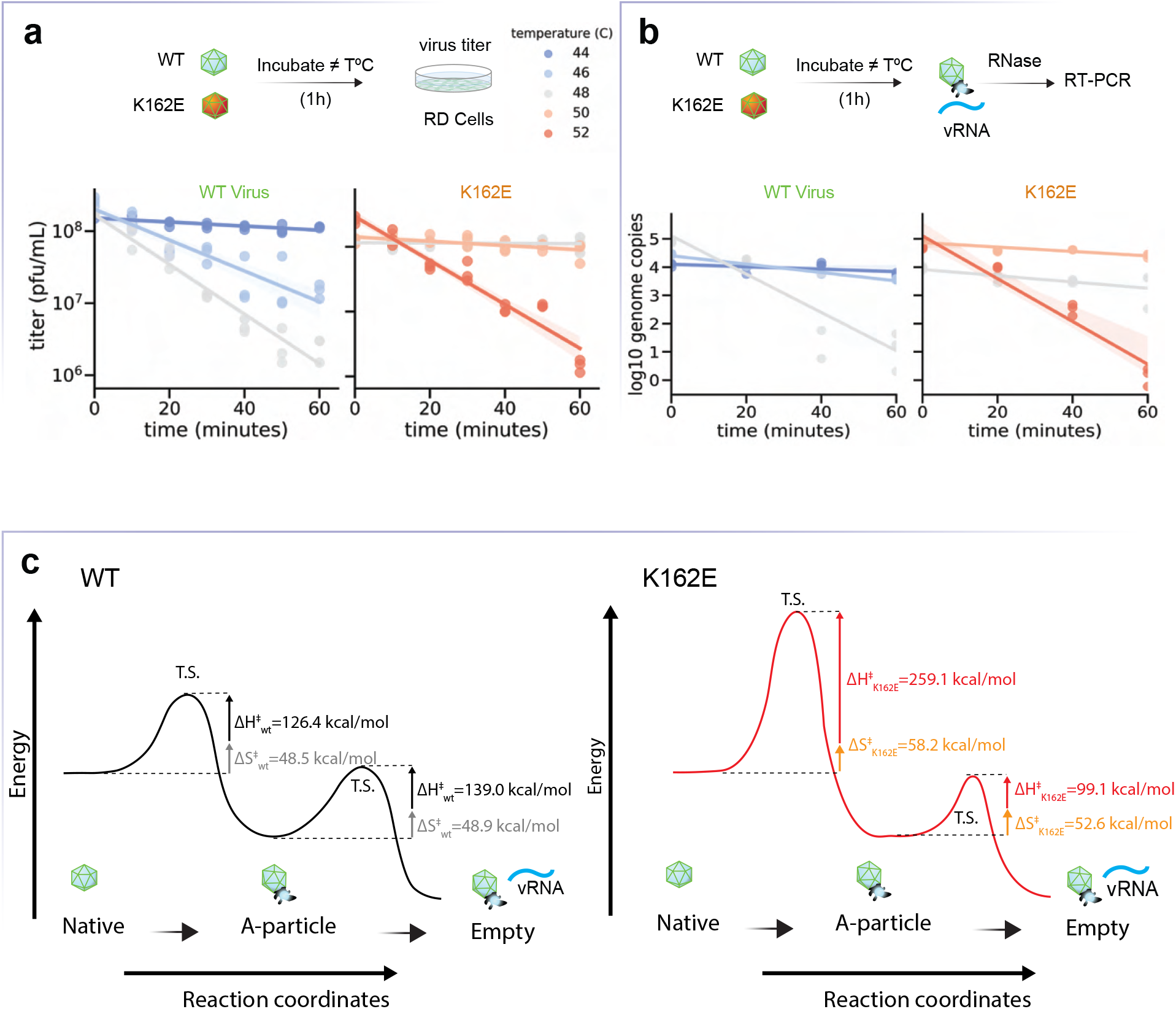
Energy landscape of uncoating state transitions. **a,b**, WT and K162E were incubated at different temperatures for 60 minutes. At the indicated time points, virus titers were determined by plaque assay (a) and proportion of viral RNA release determined by RNase-sensitivity assay (b). Model of the reaction pathway for the N-to-A and A-to-E transitions **c**, Reaction pathway for thermally mediated conversion for WT and K162E. In this model, we assume that the pathway proceeds through a single transition state (T.S.) whose free energy of activation ΔG^‡^ is composed by entropy of activation ΔS^‡^ and enthalpy of activation ΔH^‡^, which were estimated using the Eyring model. The horizontal dashed lines represent the energy barrier for the uncatalyzed reaction. Size of the arrow indicates ΔS^‡^ and ΔH^‡^ calculated for each reaction, and values of each component of the reaction are given in kcal/mol.

Enterovirus particle inactivation is accompanied by release of viral genomic RNA (vRNA). To further characterize the thermodynamics of vRNA release, we compared the rate at which WT and K162E vRNA becomes sensitive to RNase treatment after incubation at high temperatures (Fig. 3b). Genome release from K162E particles was only observed upon incubation at 52°C or higher temperatures. This contrasted with the rate of vRNA release from the WT particle, which revealed a significant increase in vRNA sensitivity to RNase at 48°C.

The conformational changes leading to WT EV-A71 virus entry and initiation of infection follow a pathway in which infectious native (N) particles are first converted to inactive altered-particles (A), which results in VP4 release and VP1 N-terminal surface exposure. To determine whether the K162E thermostable variant undergoes similar conformational transitions observed for WT virus^36^, we conducted western blotting analyses of samples taken every 10 minutes during heat treatment at 44–48°C for WT and 48–52°C for K162E. Samples were digested with trypsin prior to loading samples on the gel, which selectively cleaves only released VP4 proteins. Loss of VP4 correlated with the virus inactivation kinetics. While VP4 is completely lost at 48°C from the WT particle, the K162E retains most of the VP4 at 50°C (Supplementary Fig. 2).

Next, we used the rates of inactivation and of genome ejection to calculate the enthalpy (ΔH^‡^), entropy (ΔS^‡^) and free energy (ΔG^‡^) of the state transitions between N-A and A-E of both WT and K162E (Fig. 3C and Supplementary Fig. 3). Differently from what has been done by Tsang et al.^45^ we estimate the (ΔH^‡^) and ΔS^‡^ we took advantage of the Eyring equation:

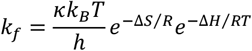

where *k_B_* is Boltzmann’s constant, *h* is Planck’s constant, and *R* is the ideal gas constant. *κ* is the transmission coefficient, set as 1, as the loss of VP4 for inactivation and genome uncoating are irreversible processes. The ΔH^‡^ of the inactivation transition from the thermostable mutant was more than twice that of the WT (126.4 vs 259.1 kcal/mol). We detected no significant change in entropy of activation for the N to A transition. Consequently, the free energy for the transition also nearly doubles, resulting in a ΔG^†^ of 235 kcal/mol. Altogether, this result suggests a change in either volume or pressure of the thermostable mutant. This increase in energy is a significant barrier for state transition considering that of the WT. Surprisingly, the ΔH^‡^ of transition for K162E was lower (99.1 kcal/mol) than for WT (139.0 kcal/mol). K162E reduces viral particle transition rate from N to A, but facilitates transition from A to E. This was mirrored in the decrease in free energy for genome ejection, with a 38.8 kcal/mol reduction in barrier-free energy. These results indicate that the mutation that was selected reduced the barrier for genome ejection to compensate for the increased barrier for priming by entering the A-particle state. There still was a net increase in the ΔH^‡^ for the complete process, where both empty complete state transitions end with the empty particle. The amplitude of both WT and K162E free energy barriers and net △H provides context for the K162E mutation inhibiting inactivation but a slight compensatory mechanism of increased genome release.

### Structural comparison of WT and K162E particles

To understand how structural changes during the uncoating process are altered by the K162E mutation, purified capsid of WT and K162E were examined by cryogenic electron microscopy (cryo-EM). Reconstructions of each particle were performed using 2D class averaging and icosahedral symmetry-based 3D reconstruction (Supplementary Fig. 4). The native state of the WT particle was refined to a resolution of 3.3Å using the gold-standard Fourier shell correlation cutoff of 0.143 (Fig. 4a, Supplementary Fig. 5). The reconstructed WT particle had a high degree of cross-correlation (score of 0.883) when compared to the reported EV-A71 native structure (EMDB ID: 20766)^46^. Enteroviruses have a deep surface depression (“canyon”) comprising the 12 pentameric vertices (Fig. 4a, light green color around fivefold axis of symmetry). The canyon is a major site of receptor binding.

**Fig. 4.**
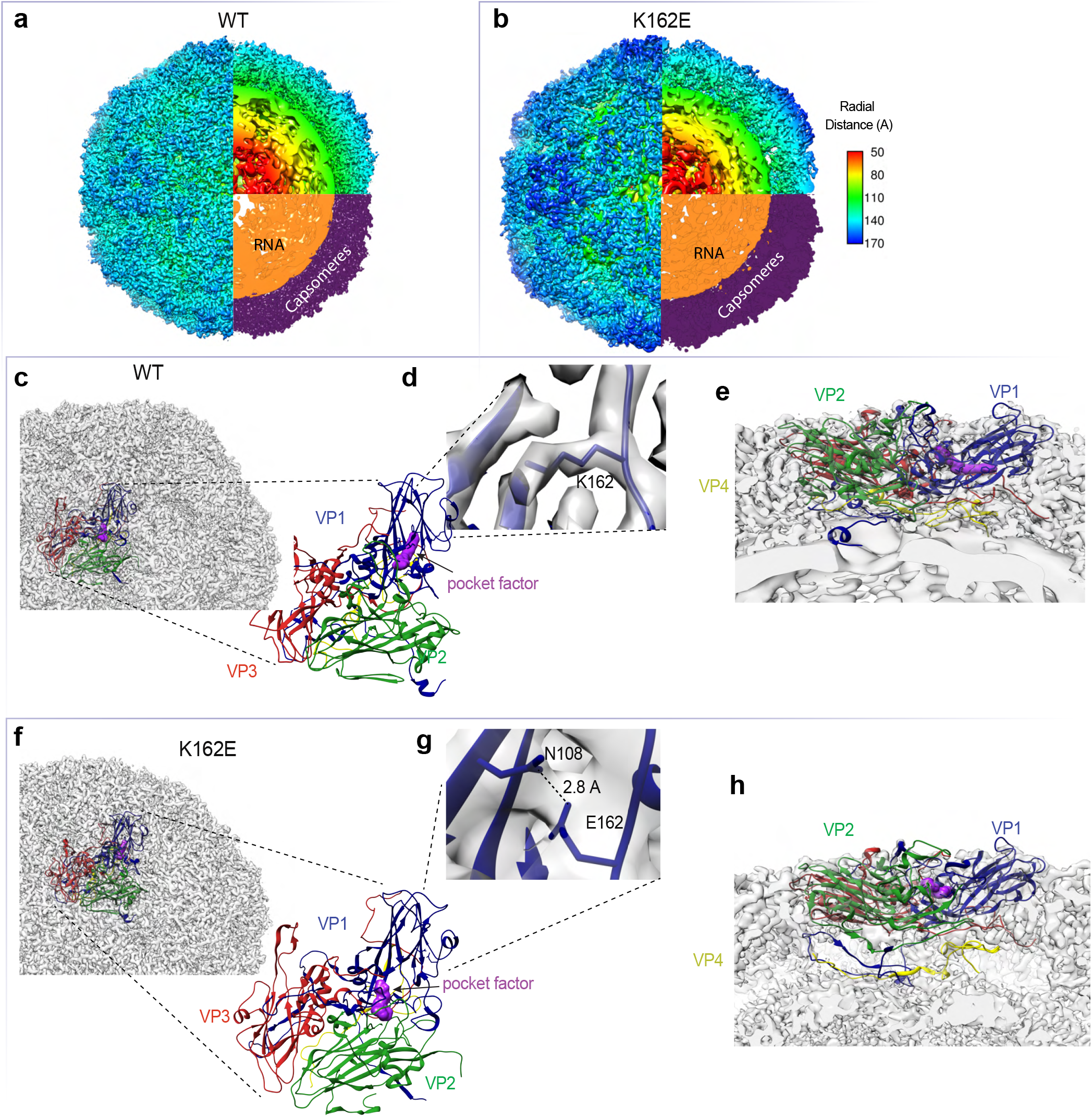
Structural determination of WT and K162E particles. **a**, WT native structure, with cross-section view, both colored radially or by RNA (orange) and protein capsomere (purple). **b**, K162E native structure, colored as described in (a). **c**, WT capsomere in context of capsid. **d**, Coulomb density of K162, indicating that the WT site of mutation does not interact with nearby residues. **e**. Side view of capsomere, showing that the N-termini of VP1 (blue) and VP4 (yellow), and the pocket factor (purple). **f**, K162E capsomere in context of capsid. **g**, K162E mutation with hydrogen-bonding residues E163-N108. **h**, Side view of K162E capsomere, showing pocket factor covered by VP1 and VP2 loops (blue and green) and VP4 (yellow) loosely associated to capsid protein shell.

We then compared the capsid structure of WT virus with that of the thermostable variant K162E. The single VP1 mutation of lysine 162 to glutamic acid drastically altered the structure of the native capsid (Fig. 4b). The particle was reconstructed to a resolution of 4.2Å. Strikingly, the K162E particle resembled an A-particle conformation of the WT particle^38^. The cross-correlation between WT A-particle map (EMDB ID: 5466)^36^ and K162E N-particle was calculated around 0.841. The thermostable K162E capsid had expanded from 30.4–31.0 nm in diameter. The K162E global capsid expansion was the consequence of both a swelled capsid protein shell and an increase in the volume of the internal cavity that was occupied by the virion RNA. This result is in line with the observed change in enthalpy.

Packaged virion RNA structures were determined independently of the outer capsid protein by masking the outer shell of the capsid during 3D reconstruction. In both WT and K162E particles, multiple concentric layers of RNA were observed at a resolution ranging from 6–14Å (Figures 4a and b, proteins in purple and RNA in orange). However, the outer layer of the K162E packaged RNA was observed to be expanded from 140 Å of the WT to 180 Å in K162E, due to the conformational change of the protein shell in K162E. The particle structure also revealed noticeable discontinuity between the RNA-capsid interface in K162E (Fig. 4b, gaps). Thus, it appears that the virion RNA expands to fill the larger volume of the internal cavity within the K162E capsid.

We next used the obtained cryo-EM maps for WT and K162E to fit an atomic model of the asymmetric capsomere that comprised one copy of each capsid protein (VP1, VP2, VP3, and VP4). The capsomere model for WT has a good cross-correlation (score = 0.883) with a previously proposed structure of the EV-A71 capsomere^37^. We observed classic “jelly-roll” beta-barrels formed by VP1, VP2, and VP3. Within that structure, we observed electron density corresponding to tightly packed lipid moiety, known as “pocket factor”. The pocket factor in the WT particle interacts directly with the VP1 beta-barrel. Lysine 162 in VP1, well resolved in the density map, does not appear to interact with nearby residues. Instead, the positively charged side chain is exposed at the surface of the particle, near the exit site of the pocket factor (Fig. 4d). The exit site of the pocket factor is near K162, but the pocket-factor and K162 do not directly interact. A side view of the WT structure reveals a tight interaction of the extended VP4 with the internal surface of the protomer and the external surface of the RNA (Fig. 4e, VP4 in yellow and RNA in orange). Similarly, the N-terminus of VP1 (blue) also extends toward the RNA.

The K162E capsid structure was also used to determine the atomic model of the asymmetric capsomere (Fig. 4f). The K162E particle maintained key structural elements in the WT structure, where the jelly-roll beta-barrel central core is nearly identical to that of WT. However, the electron density map suggested that the substituted glutamic acid at position 162 forms a hydrogen bond with nearby asparagine 108 at the VP1 beta-barrel central core, which might lead to stabilization of the VP2 EF loop (residues 132–146). The new conformation of the VP1 GH loop (residues 204–224) and VP2 EF loop appears to block the exit site of the pocket factor. Preventing pocket factor release may delay conformational changes required during the uncoating process (as described in Fig. 2e). At the interface of the protein shell and the virion RNA, most of VP4 appears to be removed from its interaction with the protomer; however, it maintains its close contact with the N-termini of VP1 and viral RNA (Fig. 4h). The portions of the VP1 N-terminus in contact with the RNA were characterized by a high content of positively charged lysines and arginines that likely form electrostatic interactions with the negatively charged RNA.

### Discrete heated particle conformations reveal details of the uncoating steps

We next studied particle conversion using purified WT and K162E particles. As described above, particle conversion from N to empty capsids (E) follows a predetermined protein unfolding process that can be mimicked by incubating the purified N particles at high temperature. Accordingly, native particles of the WT and K162E were heated at 52°C for 1 hour to convert all particles to inactivated A-particle or empty E particles. Heat-inactivated particles were reconstructed using 2D class averaging and icosahedral symmetry-based 3D reconstruction. Particles then were classified according to whether they had Coulomb density at the center of the particle. A-particles showed density, but empty particles did not (Fig. 5a and b). Atomic models were built for the native and A-particle states of WT and K162E using the asymmetric unit of the entire capsid (see Supplementary Fig. 4).

**Fig. 5.**
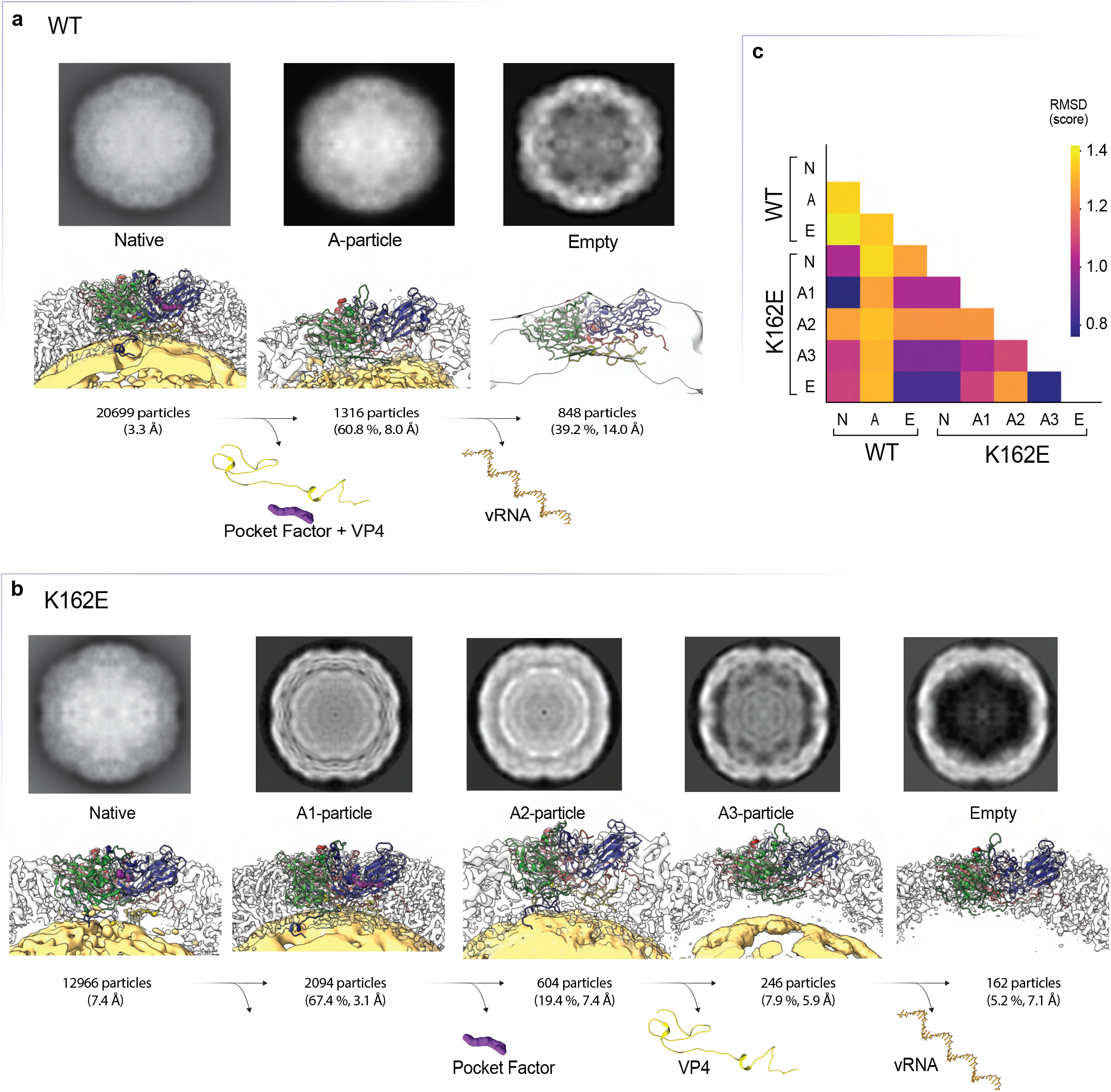
Structural determination of WT and K162E intermediates states during thermal conversion. **a**, Progression of uncoating of WT EV-A71, illustrated by 2D projection of the capsid 3D-class average and side view of the capsomere in the Coulomb density map for the native, A-particle, and empty states. Numbers of particles used for each reconstruction and resolution of the structure, as determined by the gold standard FSC 0.143 cutoff, are included below the capsid side view. Sideview also includes the atomic model built for each capsid structure. Both A-particle and empty states were reconstructed from the same heated dataset, with percentage of dataset being classified as either A-particle or empty included with number of particles and structure resolution. **b,** Uncoating states of K162E, with data presented as in (a). Extra states A1-, A2-, and A3-particles, in place of the A-particle, isolated by 3D class averaging, also presented in the uncoating pathway. The intermediates during the uncoating process irreversibly lose both the pocket factor and then VP4 prior to genome release, as indicated by lost Coulomb density. **c**, Cross correlation between capsid Coulomb-density maps of WT and K162E uncoating states. Level of cross-corre-lation is indicated by heat map representing RMSD values.

The WT A- and E-particles were resolved to a resolution of 8.0 and 14Å, respectively (Fig. 5a). Even at this relatively low resolution, the A-particle matches previously described structures (EMDB ID: 5465), with a cross-correlation score of 0.918. The empty state WT map also matched a previously solved structure (EMDB ID: 5466) with a cross-correlation score of 0.954. Despite the large difference observed in the native capsids of WT and K162E, we determined a moderate root-mean squared deviation (RMSD) score of 1.01Å (Fig. 5b).

The population of K162E particles containing Coulomb density at the center of the particle (A-particles) was heterogeneous in nature and composed of at least three distinct structures. The three states (A_1_, A_2_, A_3_) were resolved by 3D class averaging and refinement steps. The structural dissimilarity between the particles indicates three conformationally distinct states occurring during the inactivation. The proportions of A_1_, A_2_, and A_3_ in the heat-inactivated preparation were 67.4%, 19.4%, and 7.9%, respectively. These structures were resolved at 3.1, 7.4, and 5.9Å resolution for A_1_, A_2_, and A_3_, respectively. Comparing the intermediary states of both WT and K162E capsomere atomic structures by RMSD values revealed that the K162E A1 state was more like the WT native state (RMSD of 0.76Å) than the WT A-particle state (RMSD of 1.28Å). This indicates that the morphology found in the A_1_ state and the native state may be required prior to pocket factor loss. In contrast, while the comparison of WT and K162E N-particle state yields a lower RMSD score of 1.02 Å than the K162E N-particle with other WT states, the WT N-particle is more like the K162E A_1_ state.

Density maps were used to determine the presence of both VP4 and the pocket factor. Interestingly, while the most abundant state A_1_ contained both VP4 and the pocket factor, A_2_ had lost the pocket factor but retained VP4, and A_3_ no longer contained pocket factor or VP4. A trajectory of irreversible molecule loss was resolved from the presence of either none, one, or both molecules. These data suggest that the pocket factor must be dislodged prior to VP4 loss, and both molecules must be released prior to complete genome uncoating^19^. This may be due to the pocket factor sterically “locking” the capsid in a higher-energy N-particle state, compressing the viral RNA^47^. Intriguingly, RNA packing may play an energetic role during the uncoating process, where the amount of condensation of the viral RNA by its secondary structure may exert energetic pressure on the capsid protein shell^48^. The particle structure also revealed noticeable discontinuity between the RNA-capsid interface in K162E. This may be due to the tightly packed RNA expanding to fill the larger volume of the K162E capsid and releasing the tension of the “entropic spring” of condensed RNA.

### Examining the dynamics of thermostable K162E and WT capsid proteins

The structures reconstructed from our cryo-EM data clearly suggest that, during uncoating EV-A71, the particle undergoes several irreversible steps that delineates discrete stages during the entry process. However, these structural data provide only a static picture of each of the virus particles while several lines of evidence suggest that virions are dynamic structures that exist as an ensemble of reversible states^40^. To gain insights into the dynamical properties of the WT and K162E capsids, we made use of full-atom molecular dynamics (MD) simulations. Although the time and length scale accessible to MD are shorter than that of the uncoating and entry process, MD might capture differences in the dynamics of the capsid proteins underlying thermostability of EV-A71. Particularly we want to probe the local changes in the dynamics of the two viral N particles that might trigger the transition to the A particle. Given the size of the viral capsid, which is about 304Å for the WT and 310Å for the thermostable K162E, and the position of the K162E mutation, which is located on VP1 at the interface between two capsomeres (see Fig. 5b), we decided to simulate a system formed by a pentamer consisting of five capsomeres solvated in water (Supplementary Fig. 7). Such a system enables detection of possible relevant interactions between capsid proteins within the same capsomer or between adjacent capsomeres. Specifically, we first simulated the WT and the thermostable K162E native structures at 30°C for approximately 200 ns at constant number of particles N, pressure P, and temperature T, and then at 52°C. This last temperature was selected because experimentally both particles have left the N state.

We first analyzed the dynamics of the WT and K162E systems at 30°C and calculated the evolution in time of the RMSD, which is as a measure of the average deviation of a capsid protein from the initial structure at each time point of the simulation. We calculated the time evolutions of the RMSD of the capsid proteins (VP1, VP2, VP3, VP4) and sphingosine pocket factor of each capsomer for the WT and K162E pentamer (Supplementary Figs. 8 and 9). To compare the global flexibility of WT and K162E capsid proteins, we computed the RMSD mean value for each capsomer protein at each time point (Supplementary Fig. 10). The K162E RMSD values for VP1, VP2 and VP4 were significantly greater than those of WT RMSD. This suggested that, in K162E, these capsid proteins have higher flexibility than those of the WT at 30°C. In contrast, VP3 seems to be less flexible than the other proteins for both WT and K162E pentamers (Supplementary Fig. 8, 9, 10). The large size of the error bars in VP1, VP2, and VP4 also indicates higher variability in the dynamics of the single capsid proteins in K162E compared to that for WT (Supplementary Fig. 10). VP3 instead is very stable along the time evolution examined by the simulation. Interestingly, the mean RMSD of WT VP1 also shows large error bars, and at around 170 ns, it reaches the value of the K162E VP1. This is because the unstructured N-term of three VP1 capsomeres starts to unfold and flop around. Lastly, although RMSD values of the WT and K162E sphingosine are low (1 and 3Å), K162E pentamer displays greater values than those for WT and has larger error bars. This suggests that the sphingosine wiggles within the K162E VP1 pocket. This is expected given the greater mobility of the K162E capsid proteins. Next, we examined the simulations carried out at 52°C for WT and K162E pentamers. Differently form the experimental observations, the simulated WT native structure doesn’t unfold at high temperature. This is likely due to the simulation time, which might be short for this process to occur in our system. Also, the fact that we are simulating a pentamer instead of a complete viral particle might be the source of the difference with the experiments. Like simulations at 30°C, we also observed higher mean RMSD values and larger error bars for K162E than WT at the higher temperature (Supplementary Fig. 8, 10, 12).

Although the RMSD time evolution is a good metric to investigate protein dynamics, it lacks information on the local dynamics of specific loops or protein regions, which can be accessed by computing the root mean squared fluctuations (RMSFs). Therefore, to characterize the dynamics occurring in proximity of the pocket factor region and the different mobility of VP1 and VP2 capsid proteins at 30°C and 52°C, we computed the RMSF, which is a measure representing the range of amino acid position fluctuations during the simulation (Fig. 6a-b). At 30°C (Fig. 6a, top panels), the VP1 RMSFs of K162E overlap within the errors with those of the WT, whereas in the VP2 capsid protein, the RMSF values of K162E are consistently larger than those of the WT. Interestingly, the major differences between the WT and the K162E RMSF patterns were observed in the EF loop of VP2 and in the N-terminus of VP1, specifically between amino acids 30 and 70. At 52°C (Fig. 6a, bottom panels), K162E VP1 showed an increase in the dynamics of the region where the mutation is located and of the GH loop indicated higher RMSF values than the lower temperature simulation. On the contrary, the dynamics of the GH loop in the WT VP1 capsid proteins was slightly reduced. We also observed an increment in the fluctuation dynamics of the VP2 EF loop in both the WT and K162E systems. However, the large error bars in K162E indicated greater variability in the motions of the capsomeres with respect to that of the WT. Overall, upon increasing the temperature, the RMSF features remained nearly unaltered in both VP1 and VP2 of the WT-simulated system but changed remarkably in K162E relevant regions, such as in proximity of the mutation and of the GH and EF loops.

**Fig. 6.**
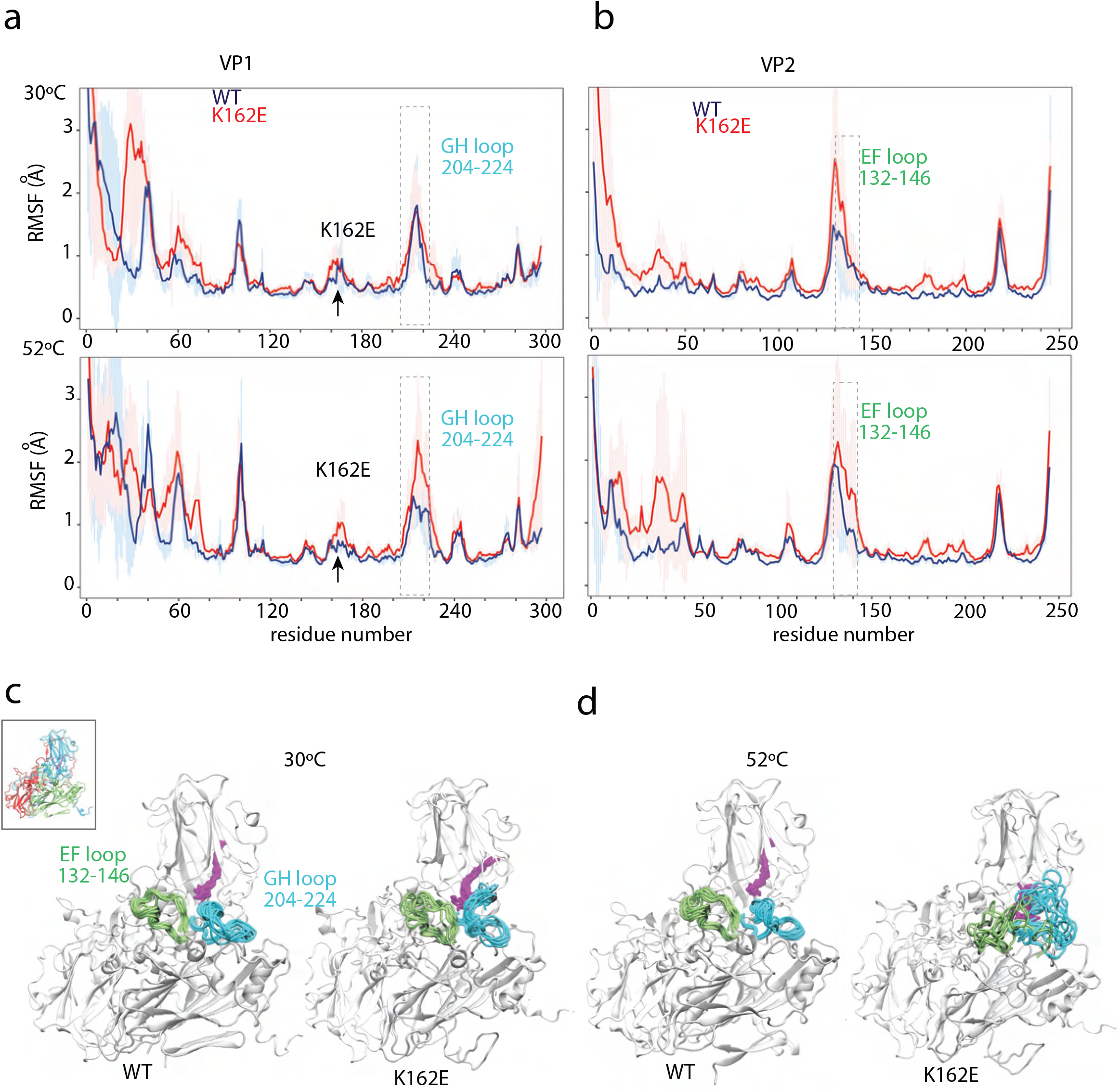
Dynamics of the EV-A71 and K162E EV-A71 pentamers at different temperatures. Representative snapshots of the EV-A71 WT (**a,b**) and of EV-A71 K162E (**c,d**) equally spaced in time over the NPT simulations at 30 (**a,c**) and 52°C (**b,d**) are overlapped to show the dynamics of the GH and EF loops of VP1 and VP2, respectively. The inset in a) shows the capsomere structure for the WT and mutant systems. The capsid proteins VP1, VP2, VP3, and VP4 are represented in white, new cartoon format, the EF loop in green, the GH loop in cyan, and the pocket factor in purple, licorice format. (**e**) VP1 and VP2 RMSFs of the WT (blue) and of the mutant (red) calculated from the simulations carried out at 30°C (top panels) and 52°C (bottom panels). The box is a guide for the eye to show the location of GH and EF loops in VP1 and VP2. The RMSFs are reported as a function of the residue number.

To gain a closer look at the dynamical trends of the EF and GH loops at different temperature revealed by the RMSF analysis, we visually inspected the simulations (Fig. 6; Supplementary Movies 1-4). Fig. 6c and d show 10 representative WT and K162E configurations of the loops and of the sphingosine extracted from different frames of the 30”C (Fig. 6c; Supplementary Movies 1 and 2) and 52°C (Fig. 6d; Supplementary Movies 3 and 4) simulations at regular intervals, including the first and the last frame. We represent the first frame of the whole capsomere in white and subsequent conformations in green (VP2 EF) or light blue (VP1 GH). At 30°C, the EF and GH loops of the WT simulation nearly overlapped along the simulation time points (Fig. 6c; Supplementary Movies 1),. Furthermore, WT EF and GH loop dynamics were not affected by increasing the temperature (Fig. 6d; Supplementary Movie 3). Compared to the WT dynamics, the K162E loops had higher mobility already at 30°C, where the loops were able to explore a large region in space (Fig. 6c; Supplementary Movies 2). At high temperatures, the K162E EF and GH loops and the proximal region, including the pocket factor, undergo dramatic conformational changes (Fig. 6d; Supplementary Movie 4).

We hypothesized that the major mobility of the K162E system with respect to the WT system could be due to the different number of hydrogen bonds (H-bonds) formed between the subunit protein pairs. Therefore, we calculated the average H-bond number at VP1–VP2 and VP1–VP3 belonging to the same capsomeres and at VP1–VP3 belonging to adjacent capsomeres (Supplementary Fig. 12). Overall, irrespective of the temperature, we observed a lower number of H-bond interactions among capsomers in K162E than the WT system, with the only exception of VP1-VP3 interactions at 52°C, where K162E and WT exhibited similar mean H-bond count within the errors. Interestingly, upon increasing the temperature, the WT had a decrease in the mean H-bond number formed between proteins belonging to the same capsomer, whereas in the K162E system, that number remained practically unaltered. On the contrary, increasing the temperature did not affect the H-bond interactions between VP1 and VP3 of adjoining capsomer. These results further indicate that the K162E structure is more flexible than WT structure and such flexibility in K162E is likely due to a reduced overall electrostatic hydrogen-bond network among capsid proteins within the same protomers.

### *In vivo* characterization of the MP4-K162E mutant virus

In cell culture, K162E has a defect in replication due to a delay in the uncoating process. Surprisingly, K162E increases virulence of EV-A71 in a mouse model of infection. We engineered a K162E mutation into MP4, a mouse adapted strain of EV-A71, and inoculated it into susceptible mice expressing the human EV-A71 receptor (hSCARB2)^49^. After intraperitoneal inoculation, 66.7% of mice inoculated with WT survived infection, compared to only 14.3% of those infected with K162E (Fig.7a, left). In addition, after intracerebral inoculation, the median survival time for the K162E variant was shorter than WT (Fig.7a, right).

**Fig. 7.**
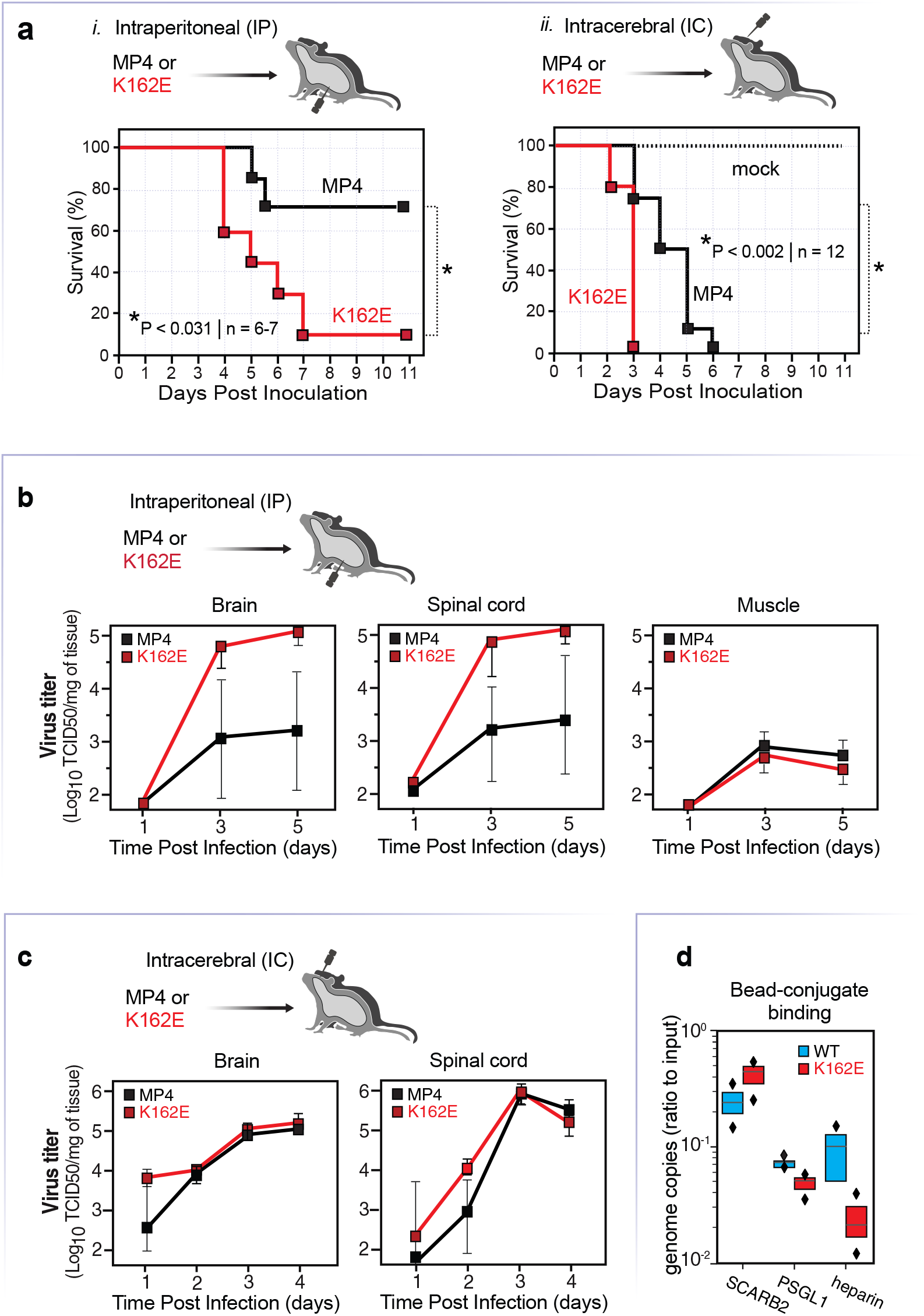
Virulence of WT (MP4) and MP4-K162E variant in hSCARB2 transgenic mouse model. **a,** Susceptibility of hSCARBE2 transgenic mice to EV-A71 K162E variant. Four-week-old mice were intraperitoneally inoculated with 1×10^5^ TCID_50_ of MP4 (shown as black) or MP4-K162E (shown as red) variant and 3-week-old mice were inoculated with 5×10^4^ TCID_50_ of the viruses and monitored for survival for 14 days. Mice in the control group were injected with the viral medium. **b**, Tissue distribution of the virus. Four-week-old mice were inoculated with 1 × 10^5^ TCID_50_ of MP4 or MP4-K162E or **c**, 3-week-old mice were intracranially inoculated with 5 × 10^4^ TCID_50_ of the viruses. Selected tissues were harvested on the indicated days for virus titration. Virus titers were determined by a standard TCID_50_ assay and shown as log TCID_50_ per gram of tissue. Results are shown as mean ± S.E.M. **d**. Beads conjugated with human SCARBE2, human PSGL1 or heparin sulfate were incubated with EV- A71 WT and K162E for 1 hour at 4°C, and then precipitated, and washed with cold PBS. Virus RNA associated with beads was determined by RT-PCR. K162E demonstrated a reduced affinity for heparin sulfate.

To determine tissue tropism of thermostable variant K162E, after intraperitoneal inoculation, we determined virus titers at 1-, 3-, and 5-days post-infection in the brain, spinal cord, and muscle tissues. (Fig. 7b). The most striking differences were found in the brain and spinal cord on days 3 and 5, where the K162E variant achieved titers that are 100-fold higher than WT. The dramatic increase in replication in neuronal tissues was not observed in muscle tissue. This suggests that K162E has an increased capacity to invade and/or replicate in the CNS.

Next, we determined virus titers in the brain and spinal cord after a direct intracranial inoculation. Higher virus titers of the M162E mutant than WT were detected in the brain on day 1 post-inoculation and in the spinal cord on days 1 and 3. (Fig. 7c). This suggests that the K162E variant has an increased capacity to enter and spread within the CNS.

To investigate if the structural change to the K162E capsid resulted in lowered immune detection, a neutralization assay was performed using blood collected from mice previously inoculated with MP4. Serum from two different mice showed comparable neutralizing titers against the WT and K162E mutant virus, suggesting that introduction of the mutation K162E should not differ the immune response in the form of neutralization antibodies (Supplementary Fig. 12).

Given the large conformational changes observed in the K162E particle and the unexpected observation that K162E is more virulent than WT, we next determined the binding efficiency of K162E to cellular receptors. We used magnetic G-beads conjugated to either shSCARB2, soluble or human PSGL1 (shPSGL1), and we measured the number of particles captured by extracting genomic virion RNA from capsids and quantified by RT-qPCR (Fig. 7d). Binding to shSCARB2 was increased by the K162E mutation, and binding to (shPSGL1) or heparin was reduced. The largest decreases in binding were found for heparin, which might be expected as the side-chain charge changes of K162E from positive to negative may have a lowered electrostatic attraction to the highly negative charge of heparin. These results suggest a possible mechanism underlying the increase of K162E in neurovirulence^50^. It was recently proposed that increased attachment to SCARB2 and a decrease of interaction with heparin sulfate is associated with an increased EV-A71 neurovirulence^51,50,52,53^. Thus, the isolated thermostable virus is more pathogenic regardless of the fact that in cell culture demonstrated reduced entry efficiency.

## Discussion

In this study, we experimentally examined the consequences of increase virus particle thermal stability. We isolated a variant carrying a single point mutation in capsid protein VP1 (Fig. 1), which displays a significant increase in tolerance to inactivation by temperature (Fig. 1 and 2). This variant is less effective to initiate infection, it is more sensitive to inhibitors of endosome acidification (Fig. 2) and has a reduced particle specific infectivity (Supplementary Fig. 2b). Structural analyses revealed a dramatic conformational rearrangement of capsid protomers, resulting in an expanded particle with an increase in volume of 6% compared to the WT virus (Fig. 4). Both capsid protein shell and virion RNA are expanded providing additional space for molecules to wiggle within the structure. This results in a lower number of H-bonds and an overall increase in the flexibility of the particle (Fig. 6 and Supplementary Fig. 7-14). These results are surprising. As the K162E particle is significantly more thermotolerant (Figs. 1 and 2) and hyper-dependent to lower endosomal pH (Fig. 2e), the expectation was that additional intramolecular interactions should result in a more stable particle so that it should better tolerate an increase in temperature. Our results provide an alternative view to how viruses modulate particle stability, while preserving infectivity.

Our study suggests that virus particle stability can be regulated by modulating the intrinsic energy of the particle. It is thought that both enveloped and non-enveloped viruses rely on energy accumulated in their structural proteins to disrupt cell membranes to deliver the virus genome into the cytoplasm of the cell and initiate infection^54^. Accordingly, virus particles are like “spring loaded” structures^55^, in which accumulated energy can be released upon encountering appropriate conditions, such as pH, membranes, virus receptor, during entry. In the case of non-enveloped viruses, like EV-A71, it is thought that part of the energy generated during the genome synthesis can be employed for RNA packaging, which, in turn, is forced to associate with capsid proteins and form a “tense-state” RNA-protein complex: the virus particle^56,57^. The energy accumulated in the “spring loaded” particles can be then used to facilitate capsid protein conformational changes, membrane disruption, and RNA delivery into cells^58^. The results described in our study suggest that the thermotolerant K162E variant is structurally more flexible than the WT particle. Thus, the K162E relax particle may have lower intrinsic enthalpy and less energy to facilitate genome release. In this way the energy required to overcome the transition state activation, enthalpy of activation (Fig. 3), for the conversion to an A-particle is increased, which results in a decrease in virus ability to initiate infection. At the same time, a relaxed particle with lower accumulated energy and higher flexibility can withstand higher temperatures or environmental challenges, by releasing part of the energy through the thermal motion of flexible-loops^59^.

It is generally accepted that heat inactivation of enterovirus particles offers a simpler approach to examine the physiological relevant conformational change trajectories that lead to virus uncoating^15,18,38^. The unbiased 3D classification of K162E particles generated during heat inactivation (Fig. 5) provided novel details in the uncoating process that were poorly understood^60^. We observed a couple of intermediates that suggest that the first step in particle conversion is the release of the pocket factor (A1 particle to A2 particle), followed by large rearrangement of the N-terminus of VP1 and release of VP4 (A2 particle to A3 particle). Finally, virus RNA is expelled generating empty particles (A3 particle to empty particle). A key initiating event in this pathway is the release of the lipid pocket factor, which may be facilitated by the proximity of hydrophobic cell membranes and the primary EV-A71 receptor SCARB2 (Fig. 2c). This process may be akin to that observed in enveloped virus, where the hydrophobic fusion peptide undergoes a large spatial rearrangement, facilitated by the proximity of cell membranes and insertion in the cell membrane^61^. The movement of the fusion peptide is thought to drive additional changes in virus structural proteins leading to membrane fusion and virus entry. The next step, VP1 N-terminus and VP4 release, has been proposed to form an umbilical channel through the membrane that enables viral RNA translocation through the cell membrane^18^. The identification of additional intermediates during particle heat inactivation (Fig. 5) supports this previously proposed model. An additional observation in our study is that the K162E expanded particle, which resembles WT A particles (Fig. 5c), undergoes an additional step before the uncoating process described above takes place. Namely, the K162E particle shrinks to adapt a conformation similar to that observed in the native WT particle (Fig. 5c). The shrunken structure (A_1_ particle Fig. 5b) is no longer infectious, even though it is very similar to the infectious WT mature particle, because the pocket factor exit is blocked by the EF-VP2 and GH-VP1 loops, preventing release of the pocket factor and subsequent conformational changes. The effect of the VP1 and VP2 loops blocking the pocket factor release is similar to that elicit by small-molecule inhibitors that bind with high affinity into a hydrophobic pocket within VP1 limiting infectivity of many picornaviruses^19^.

Another unexpected result from this study was that the K162E variant, which is less efficient in uncoating viral RNA and initiating infection under physiological conditions, was significantly more virulent in a mouse model of infection (Fig. 7a). Indeed, K162E more effectively accessed and replicated in the central nervous system than the WT virus. While the K162E VP1 mutations do not affect virus interaction with SCARB2 and PSLG1, the variant shows a fivefold reduction in its ability to interact with heparin (Fig. 7d). We speculate that the K-to-E mutation changes the charge on the particle surface to affect the interaction with cationic heparin. Interaction with heparin has been suggested to be a cell-culture adaptation and increases pathogenesis^62,63^. However, additional studies would be necessary to fully address why K162E changes tissue tropism and neurovirulence.

Understanding the biophysical properties of the balance between these phenotypes has numerous potential benefits. Many bioengineering attempts have been made to imitate the various viral endocytosis processes to deliver pharmaceuticals, such as small-molecule compounds^64^ or mRNA vaccines^65^. However, these approaches tend to have limitations with regards to not being tuned to release their cargo easily^66^ from overstabilization or are too thermo- or acid-labile to be intact through physiological environments, such as the GI tract. Empty capsid (EC) systems are an active area of research, where the viral capsid minus the genome can be used as a type of vaccine^67,68^. Barriers to development of these systems are again due to stabilizing the viral capsid to survive outside the cell. The stabilization of the EV-A71 capsid by disrupting the uncoating pathway may be a future method of EC stabilization.

## Acknowledgments

This work was supported by NIH (R01 AI36178, AI40085, P01 AI091575), the Bill and Melinda Gates Foundation to R. A. and the DARPA Intercept program (Contract No. HR0011-17-2-0027) to R.A. and S. B. We thank members and faculty of the UCSF Cryo-EM core facility, including Daniel Asarnow, David Buckley, Glenn Gilbert, and Zanlin Yu. UCSF Cryo-EM equipment at UCSF is partially supported by NIH grants S10OD020054 and S10OD021741. We also thank the members of A.C.’s thesis committee; John Gross, Adam Frost, and Joe Bondy-Denomy. We thank Patrick Dolan, Orly Laufman, and Ranen Aviner for their suggestions on the manuscript. We acknowledge S. Wheeler for providing the sphingosine force-field. We acknowledge support from the IBM Research AI Hardware Center, and the Center for Computational Innovation at Rensselaer Polytechnic Institute for computational resources on the AiMOS Supercomputer.

## Author Contributions

A.C. conceived of this study. A.C., M.T.Y., S.C. and R.A. designed the project and experiments. A.C. prepared, collected, and analyzed cryo-EM data. M.T.Y. performed the animal experiments. S.C. designed and carried out the MD simulations. R.A., S.C. and S. B. provided guidance and support. All authors wrote the manuscript.

## Resource Availability

Data and code availability. All electron micrographs used for structural determination were deposited in both PDB and EMDB. WT native, A-particle, and empty states were deposited in the PDB as 8E2X, 8E2Y, and 8E31 and deposited into EMDB as EMDB as EMD-27850, EMD-27851, and EMD-27853 respectively. K162E native, A1-particle, A2-particle, A3-particle and empty states were deposited in the PDB as 8E38, 8E39, 8E3A, 8E3B, and 8E3C and deposited into EMDB as EMD-27859, EMD-27860, EMD-27861, EMD-27862, and EMD-27863.

## Method Details

### Cells and viruses

RD cells were obtained from ATCC (CCL-136) and maintained in Dulbecco’s Modified Eagle Medium (DMEM) supplemented with 10% fetal bovine serum and 1X penicillin-streptomycin-glutamine at 37°C at 5% CO_2_. The EV-A71 strain used in this study was the 4643 strain isolated from the 1998 epidemic in Taiwan^69^, which had been ligated onto the pUC plasmid.

### Propagation and titering of virus

Using infectious cDNA clones of enterovirus A71 (EV-A71 strain 4643/TW/98 clade C2, the carrying plasmids were linearized then used as a template for *in vitro* transcription of virion RNA. 20 μg of RNA was transfected into 4.0×10^6^ RD cells. Resulting virus was propagated in RD cells from a passage 0 (P0) stock to form the initial passage 1 (P1) population. All viral populations were titered by either TCID_50_ or plaque as described.

### Passaging virus under temperature selection

The initial virus passage (P1) was either kept on ice for 1 hour (control passage) or heated at 46°C for 1 hour (heated passage) before being added to an ask of 10^7^ RD cells. Virus was titered before and after heating to ensure each passage was started with 10^5^ TCID_50_ of virus for a consistent multiplicity of infection (MOI). After 24 hours, the plates were checked for cytopathic effect (CPE) and frozen at −80”C. Three cycles of freezing and thawing at room temperature were performed to lyse viruses from cells. Supernatant from each ask was centrifuged at 2500×g for 5 minutes to pellet cellular debris, and the resulting supernatant was stored at −80°C and used for subsequent passages. Viral supernatant from each passage was further amplified at a MOI of 5 in 10^7^ RD cells, allowed the viruses to replicate for 8 hours, then lysed by Trizol (ThermoFisher), and RNA was extracted by phenol-chloroform. RNA was reverse transcribed using Superscript III (ThermoFisher) and primers specific to the P1 region of viral genome. Amplified cDNA was then used for Sanger sequencing and mutations were probed for in each passage.

### Generation of mutant virus from cDNA

Using the previously mentioned infectious clone of EV-A71 4643, the plasmid was linearized and used as a template for *in vitro* transcription of virion RNA. 20 μg of virion RNA was then transfected into 4×10^6^ RD cells. Transfected virion RNA was allowed to replicate to produce viral particles.

### Virus purification

EV-A71 was propagated in 20 100-mm dishes of confluent RD cells. Upon total CPE, virus particles were lysed with 0.5% IGEPAL CA-630 and followed by three cycles of freeze-thaw. Resulting supernatant was then centrifuged for 5 minutes at 3000×g to pellet cell debris, and the resulting supernatant was precipitated with 8% final concentration PEG 8000 at 4°C for 72 hours. Virus was pelleted at 3000xg for 1 hour, re-suspended in purification buffer and remaining debris pelleted at 3000xg for 15 minutes. Supernatant was then purified through a 30% sucrose cushion centrifuged at 100,000×g for 3 hours, followed by fractionation by 15–45% sucrose gradient at 100,000×g for 3 hours. Fractions were then dialyzed by Zeba desalting column and concentrated using a Amicon 100,000 MWCO filter.

### Thermostability assay

Purified virus samples were heated using calibrated water heat bathes set for indicated temperatures. Samples were added to the heat bath simultaneously and removed at indicated timepoints and placed on ice for at least 5 minutes. Samples used for genome quantification were treated with RNase If for 30 minutes at 37°C, followed by RNA purification by phenol-chloroform extraction and precipitation by isopropanol-sodium acetate. Viral genome quantification. Genome quantification was performed by RT-qPCR using NEB’s Luna One-Step kit. Primers (GCAAACTGG-GACATAGACATAAC) and (GGACAACTTGCCCGGTAG) were used with viral samples or IVT genomic RNA for standard curves. Samples were run on a BioRad CFX Connect thermocycler. C_q_ values were used to determine viral genome copy numbers.

### Viral particle quantification

Known titers of purified native viral particles were mixed with known concentrations of 50-nm latex beads. Samples were then imaged by negative-stain electron microscopy. Electron micrographs were then used to determine the ratio of viral particles to latex beads and calculate the concentration of viral particles. Ratios of viral particle concentrations and titer were used to determine viral particle to infectious unit ratios.

### Neutral red assay

Neutral Red virus were produced by inoculating RD cells with viral stock and allowing virus to attach and enter cells over 1 hour at 37°C. Inoculum was then removed and replaced with viral medium containing 20 M neutral red. Flasks of infected cells were wrapped in foil then returned to a 37°C incubator for 24 hours. Viral supernatant was obtained as described and kept in darkness. During assaying Neutral Red virus was serially diluted and added to 6-well plates of 5×10^5^ RD cells/well in 500-μL aliquots.

For the chemical perturbation assay, RD cells were incubated with the indicated amount of compound for at least 1 hour prior to inoculation with virus. Viral inoculum also contained the indicated concentration of compound prior to inoculation and was added to 5×10^4^ cells in a 24-well plate. Virus was allowed to attach and enter cells for 1 hour at 37°C before inoculum was removed and replaced with cell culture medium containing the same concentration of indicated compound. The virus was allowed to replicate at 35°C for 8 hours before the plates were moved to a −80°C freezer to halt replication. Samples were freeze-thawed 3X before being centrifuged to remove cell debris. Resulting supernatant was then titered.

### Western blotting

Purified viral capsids were pre-treated with 10 ng of sequencing grade trypsin (Promega) prior to denaturing sample with 2X Laemeli buffer (BioRad). Samples were then heated at 94°C for 10 minutes, followed by cooling on ice for at least 2 minutes. Samples were loaded onto a 4–20% SDS-PAGE gel (BioRad) and electroporated for 1 hour before being equilibrated in transfer buffer for 5 minutes and transferred to an activated PVDF membrane at 40V for 40 minutes then 100V for 1 hour. Membranes were blocked with 3% BSA in TBST buffer overnight at 4°C with gentle rocking, followed by washing 5X with TBST and incubation at RT with primary antibodies against VP2, mAb979 (Millipore Sigma), or VP4, CF594 (Biorbyt), for 2 hours. Membranes were washed 5X with TBST buffer and incubated with secondary antibody anti-rabbit IgG-HRP or anti-rabbit IgG-HRP in TBST buffer at RT for 2 hours. Membranes were washed 5X in TBST buffer and developed using an ECL system (ThermoFisher). Images were taken using a BioRad ChemiDoc.

### Electron microscopy

Negative-stain electron microscopy was performed after fixation of purified particle with 0.01% final concentration EM-grade paraformaldehyde. Quantifoil 400-mesh formvar copper grids were glow discharged prior to the addition of sample to the gird. Samples were allowed to adhere to grids for 5 minutes, followed by wicking away of excess liquid by applying a Whatman paper to the edge of the grid. Grids were then washed with deionized distilled H_2O_, and excess liquid wicked before applying 2% neutral phosphotungstic acid for 30 seconds. Excess liquid was wicked away and grids were imaged with a FEI Tecnai T12 120 kV microscope with a Gatan UltraScan 4k CCD camera.

Cryo-EM was performed by first preparing grids using a ThermoFisher Mark IV vitrobot. 3 μL of sample was applied to a Quantifoil R 2/1 ultrathin carbon grids for 5 minutes to allow the sample to adhere to carbon before excess liquid being wicked away with Whatman paper. Grids were then placed in the vitrobot at 100% humidity and 3 μL of sample applied again for 1 minute. Excess liquid was then blotted using blot-force 3 for 17 seconds and rapidly plunge-frozen in liquid ethane. Grids were then loaded into either a Talos Arctica or Glacios 200 kV microscope with Gatan K3 detectors. Images were collected on the Arctica at a nominal magnification of 28,000X with a dose rate of 21 e^-^/pixel/second for 6 seconds for a total of 0.53 e^-^//frame. Images were collected on the Glacios at a nominal magnification of 34,000X at 16.5 e^-^/pixel/second for 6 seconds for a total of 0.56 e7/frame. Image Processing. Micrograph image stacks were dose-weighted and corrected for beam-induced local motion using MotionCorr2^70^ Images were then CTF corrected using CTFFIND4 within the Relion3.1 package^71,72^. From corrected images particles were manually picked and class averaged within Relion3.1. High-contrast class averages were used for *ab initio* structure refinement. Particles were used to produce four classes of initial maps, from which the best resolved class was used for structural refinement. One round of polishing was used to produce the final Coulomb potential density map. Maps were checked for handedness and masked using a low-resolution map.

### Atomic model building

UCSF Chimera was used to fit the initial atomic models into the Coulomb potential density maps, PDB:3VBS^16^ for WT and PDB:4N53^38^ for K162E. Asymmetric maps were boxed using a 5Å mask created from the initial model, and multiple rounds of real space refinement were performed until cross-correlation between model and map was not improved by additional rounds of refinement in Phenix^73^. Capsomeres were then used to create both the pentamer and full capsid atomic model. From the whole-capsid atomic model a 5Å mask was created and used to create from the full-capsid Coulomb-density map both the virion RNA and protein capsid components. The mask was then used with the subtract functionality in Relion3.1 to iteratively refine both the protein capsid and packaged virion RNA maps. A final round of atomic model building was used with the resulting capsid protein Coulomb-density map in Phenix. All images of atomic models and Coulomb-density maps were rendered using UCSF ChimeraX^74^.

### EV-A71 and K162E EV-A71 simulation setup

The initial coordinates of the EV-A71 native pentamer and the K162E EV-A71 native mutant pentamer were extracted from structures PDB ID 8E2X and 8E38. We used the same procedure to minimize, equilibrate, and simulate the two pentamer systems at temperatures of 30°C and 52°C, which we described in the following. Both EV-A71 and K162E EV-A71 pentamer structures are composed of five capsomeres, each of them including VP1, VP2, VP3, and VP4 capsid proteins, and the sphingosine pocket factor. In both systems, the N- and C-terminus of all proteins forming the pentamers were capped. We modeled all the His amino acids in the Nδ1 tautomeric state and maintained positive charge for Arg/Lys and negative charge for Asp/Glu. The sphingosine was model as neutral. To build the initial cell box, we placed the pentamer in the center of the Cartesian coordinates and overlapped with a water box. Water molecules overlapping with the protein atoms were removed, and potassium ions were added to maintain charge neutrality. The EV-A71 simulation system contains approximately 419,000 atoms, and the K162E approximately 417,000, respectively. All full-atom molecular dynamics simulations were performed using NAMD 2.14^75,76^ with the CHARMM36m force field for the protein an ions^77^ and the TIP3P model for water^78^. The sphingosine parameter and topology files were courtesy of S. Wheeler (see Acknowledgments). We minimized the system energy using the conjugate gradient algorithm for 8,000 steps and gradually heated the simulated cell from 25°K to 303°K (30°C) in steps of 25°K. To equilibrate the system, we applied harmonic restraints to the protein and sphingosine backbone, water, and ions, and we released them during consecutive NPT simulations (constant number of particles N, pressure P, and temperature T) of 1 ns length. After this equilibration procedure, we started the production runs. To carry out simulations at the temperature of 52°C (325 K), we extracted the last frame of the simulation at 30°C, increased the temperature of 22°C and ran a 1-ns simulation before starting the production runs. For all simulations, we used a Langevin dynamics scheme to keep the temperature constant and an anisotropic coupling in conjunction with Nosé-Hoover Langevin piston algorithm to keep the pressure constant at 1 atm^79,80^. Periodic boundary conditions were applied in three dimensions. We employed the smooth particle-mesh Ewald summation method to calculate the electrostatic interactions^81,82^, and the short-range real-space interactions were cutoff at 12Å using a switching function at 10–12Å. To integrate the equations of motion during the production runs, we used a time step of 4 fs for the long-range electrostatic forces, 2 fs for the short-range non-bonded forces, and 2 fs for the bonded forces by means of a reversible, multiple time-step algorithm^83^. The SHAKE algorithm^84^ was used. Coordinates were saved every 20 ps.

The EV-A71 pentamer simulation at 303°K (30°C) was extended for 213.22 ns and that at 325°K (52°C) for 182.08 ns. The K162E EV-A71 pentamer simulation at 303°K (30°C) was extended for 192.06 ns and that at 325 K (52°C) for 191.28 ns. Simulations were carried out partially on the IBM Artificial Intelligence Multiprocessing Optimized System (AiMOS) supercomputer and partially on a cloud bare metal machine.

The simulations were visualized using VMD software^85^. The analysis of the trajectories was carried out using MDAnalysis^86,87^ and our own codes. We first computed the RMSD of the α Carbon atoms of VP1, VP2, VP3, VP4, and sphingosine of the pentamer of both systems as a function of time separately, and then we averaged the RMSD values at each time point so that we obtained the time evolution of the mean RMSD of the α Carbon atoms of VP1, VP2, VP3, VP4, and sphingosine along with the standard deviation of the mean. The root-mean-squared-fluctuations (RMSFs) were calculating by averaging over the last 10 ns of the simulations. By following the RMSD analysis procedure, we first computed the RMSF of VP1, VP2, VP3, VP4, and sphingosine of the pentamer singularly, and then we averaged the RMSF values over the five capsomers and obtained the mean RMSF and the standard deviation of the mean. The H-bond analysis was carried out by using the HBonds VMD plugin. We defined a hydrogen when the distance between the donor (D) and the acceptor (A) was 3.5Å and the D-H-A angle was 30°. We calculated the number of H-bonds formed throughout the last 10 ns of the simulations. The data represented in Fig. Supplementary Fig. 11 are the mean values calculated over the five capsomeres and the standard deviation of the mean.

### Ethics statement

All animal experiments were conducted in accordance with the guidelines of Laboratory Animal Center of National Institutes of Health. The Institutional Animal Care and Use Committee of University of California, San Francisco approved all animal protocols (Approved protocol number AN194006-01A).

### Animals

C57BL/6 mouse strain with the hSCARB2 transgene^88^ was obtained from Dr. Satoshi Koike (Tokyo Metropolitan Institute of Medical Science, Tokyo, Japan). Mice were bred in-house and maintained in a 12/12 light cycle with standard chow diet under specific-pathogen free condition in the AAALAC-certified animal facility at UCSF. Both male and female, 3–4-week-old mice were used in this study.

### Infection of mice

Mice were injected intra-peritoneally (i.p.) with 100 μl of inoculum delivering 1×10^5^–1×10^6^ TCID_50_ of virus per mouse or intracranially with 5 μl of inoculum delivering 5×10^4^ TCID_50_ of virus. Survival of inoculated mice were monitored for 14 days. Mice with the appearance of paralysis on both posterior limbs, the humane endpoint, were euthanized. For the virus tissue-distribution assay, three mice from each virus group were euthanized, perfused with 1X PBS, and selected tissues were collected aseptically, weighted and stored at −80°C. Tissue samples were homogenized in viral medium, disrupted by three freeze-thaw cycles and cleared at 1500×*g* for 10 minutes at 4°C. Virus titer in cleared supernatant was determined by a standard TCID_50_ assay.

### Neutralization assay

Serum was collected from mice inoculated with serum serially diluted twofold in medium and incubated with 100 TCID_50_ of either WT or K162E at 37°C for 1 hour. Viral samples were then added to RD cells and allowed to replicate for 7 days where total CPE was observed in infected cells. The limiting dilution factor was used to report the efficacy of serum. Serum was used from two independent mice and titers were taken in triplicate.

### Statistical analysis

All analysis was performed using Python3.7 using the Scipy.Stats module. Data was represented as mean and standard deviations of technical triplicates, with biological duplicates performed when indicated. Kinetics analysis was performed using log-normalized fitting of a linear curve using the least-squares method where the slope of the line was used to determine respective rates. Survival curves were analyzed using a Cox regression model. Student’s *t*-test was performed to compare virus titers at collected time points.

