## Supplemental Files for "A tradeoff between enterovirus A71 particle stability and cell entry"

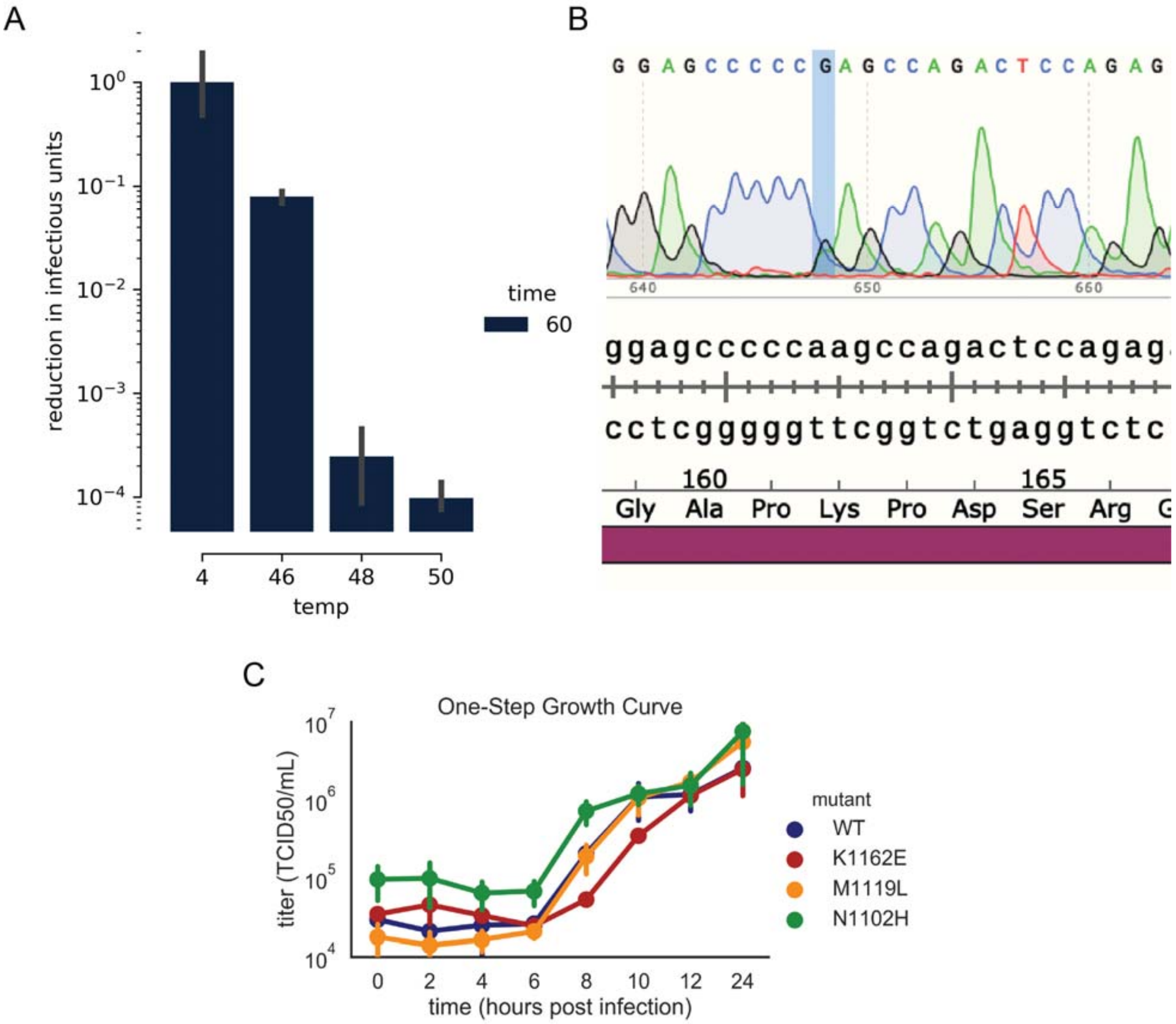


**Fig. 1.** Optimization of temperature selection, identification and characterization of thermostable variants. A) Optimization of temperature selection. Wild-type (WT) EV-A71 was subjected to different temperatures for 1 hour. The degree of virus inactivation was determined by tissue-culture infectious dose 50 (TCID$_{50}$) in RD cells. B) Sequencing analysis of heat-selected virus at passage 6. A->G transition observed at position 2924. C) One-step growth curve of viruses carrying mutations identified in treated populations.

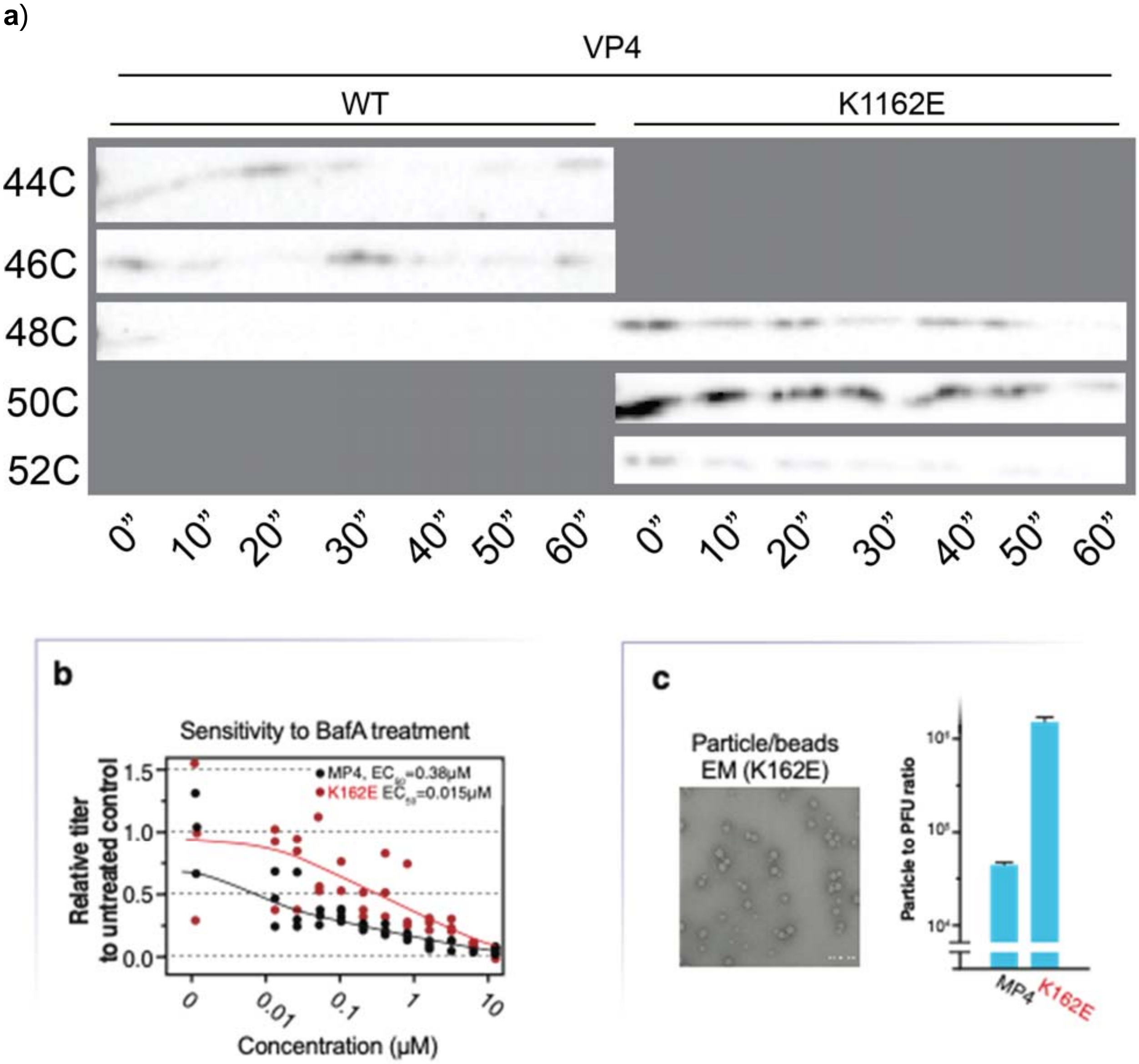


**Fig. 2.** Particle characteristics. **a**) Examining the presence of VP4 in WT and K162E particles. Purified virus particles were heated at different temperatures, and samples were obtained at 0, 10, 20, 30 , 40, 50 and 60 minutes, digested with trypsin for 30 minutes, and examined for the presence of VP4 by western blotting. **b**) To determine effective concentration of BafA1 required to inhibit viral entry, we used different concentrations of BfaA1 in RD cells prior to WT and K162E infection. BafA1 was added at least 1 hour prior, and cells washed prior to the addition of virus. Infected cells were allowed to replicate virus until 8 hours post-infection where the number of successful replications was determined by $TCID_{50}$. The EC50 of inactivation was found to be 0.38 M for WT and 0.015 M for K162E, representing approximately a 20-fold increase in inhibition. **c**) To determine if the number of physical particles required for a successful infection was also affected, the specific infectivity viral particles (physical particles to

infectious units) was calculated. The concentration of physical particles was measured against a 50-nm latex bead standard of known concentration, and the concentration of infectious particles was measured by plaque assay. The number of physical units required to initiate a successful infection, in cell culture, was increased about 20-fold by the introduction of K162E.

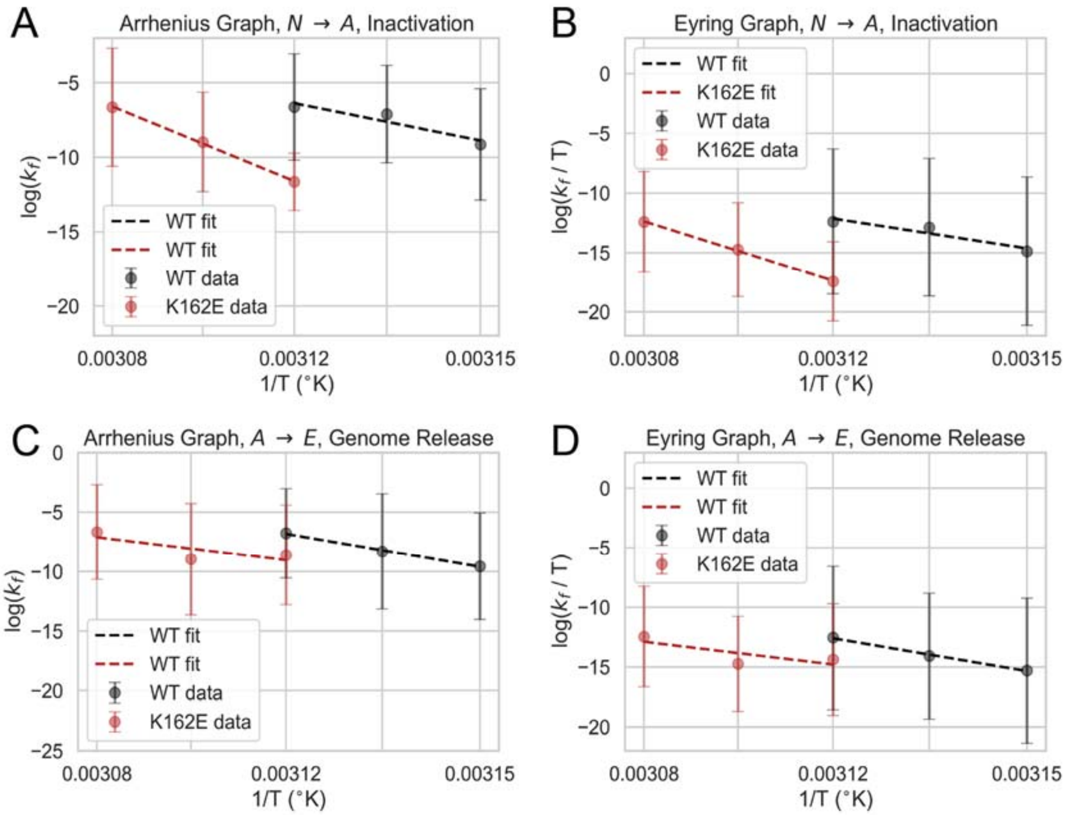


**Fig. 3**. Arrhenius and Eyring plots of inactivation and uncoating. A) Log-normalized rates of inactivation of WT and K162E from 44º to 52ºC. Rates were curve-fit from data in Figs. 3a and 3b using log-normalized least squares. The relationship between log-normalized inactivation rates and the inverse of temperature was fitted by log-normalized least-squares to determine the slope of the curve, yielding the activation energy ($E_a$). B) Log-normalized rates of inactivation for WT and K162E over the respective temperatures. Rates were determined as mentioned in SM-3A. Intercept of fitted curve was used to determine the enthalpy (**Δ**H$^{\dagger}$) from the slope of the curve and the entropy (**Δ**S$^{\dagger}$) from the intercept of the curve. C) Log-normalized rates of genome release of WT and K162E from 44º to 52ºC. Rates and curve-fitting was performed as described in SM-3A. D) Log-normalized rates of genome release for WT and K162E over the respective temperatures. Rates and curve-fitting was performed as described in SM-3.

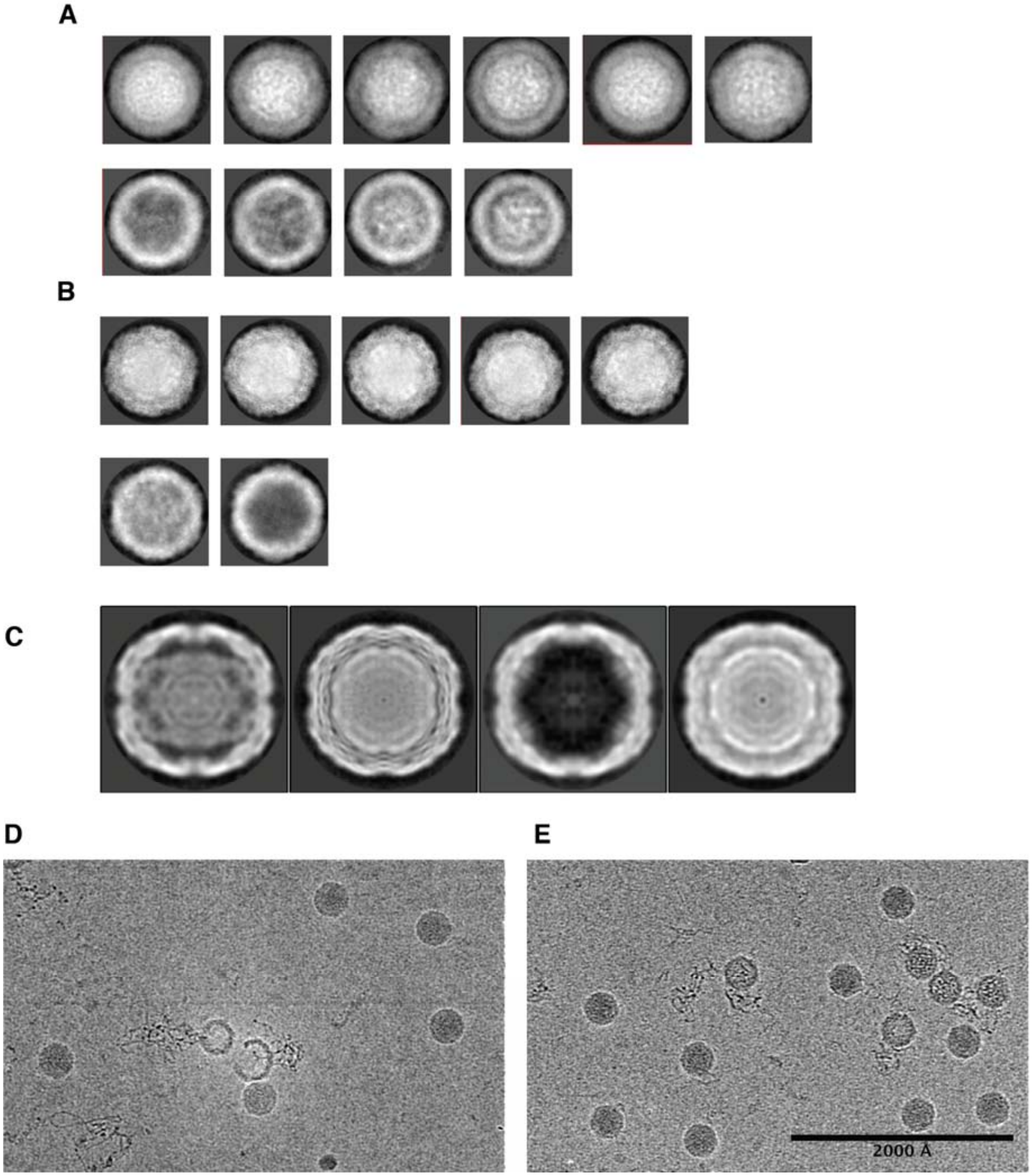


**Fig. 4**. Images of heated WT and K162E particles. A) 2D class averages of heated WT particles, either used for A-particle (top row) or empty particle (bottom row) reconstruction. B) 2D class averages of heated K162E particles, either used for pre-A, A-particle, or post-A (top row) or empty particle (bottom row) reconstructions. C) Projections of 3D class averages of heated K162E particles. From left to right, projections are of empty, A-particle, post-A, and pre-A states. D) Sample micrograph of heated WT particles. E) Sample micrograph of heated K162E particles.

| | WT-native | WT-A-particle | WT-empty | K162E-native | K162E-$A_1$-particle | K162E-$A_2$-particle | K162E-$A_3$-particle | K162E-empty |
|---|---|---|---|---|---|---|---|---|
| # particles | 20699 | 1316 | 848 | 12966 | 2094 | 604 | 246 | 162 |
| Map resolution (A) | 3.3 | 8 | 14 | 4.1 | 3.1 | 7.4 | 5.9 | 7.1 |
| Detector pixel size (A) | 0.72 | 0.72 | 0.72 | 0.603 | 0.72 | 0.72 | 0.72 | 0.7 |
| Nominal magnification | 28000 | 28000 | 28000 | 34000 | 28000 | 28000 | 28000 | 28000 |
| Electron exposure ($e^-/A^2$) | 62 | 62 | 62 | 68 | 62 | 62 | 62 | 62 |
| Voltage (kV) | 200 | 200 | 200 | 200 | 200 | 200 | 200 | 200 |
| **Atomic Model** | | | | | | | | |
| Cross-Correlation | 0.85 | 0.62 | | 0.74 | 0.83 | 0.64 | 0.71 | 0.61 |
| MolProbity Score | 1.44 | 2.01 | | 2.99 | 1.51 | 2.19 | 1.73 | 2.46 |
| Clash Score | 6.43 | 11.06 | | 10.93 | 4.49 | 10.39 | 7.70 | 15.57 |
| Poor Rotamers | 0.14 | 0.17 | | 8.41 | 0 | 0 | 0 | 8.35 |
| Favored Ramachandran | 97.6 | 92.95 | | 79.86 | 96.04 | 91.73 | 95.5 | 96.55 |
| Allowed Ramachandran | 2.4 | 6.9 | | 19.42 | 3.96 | 8.27 | 4.5 | 3.45 |
| Disallowed Ramachandran | 0 | 0.15 | | 0.72 | 0 | 0 | 0 | 0 |

**Fig. 5.** Summary statistics of cryo-EM and atomic model building

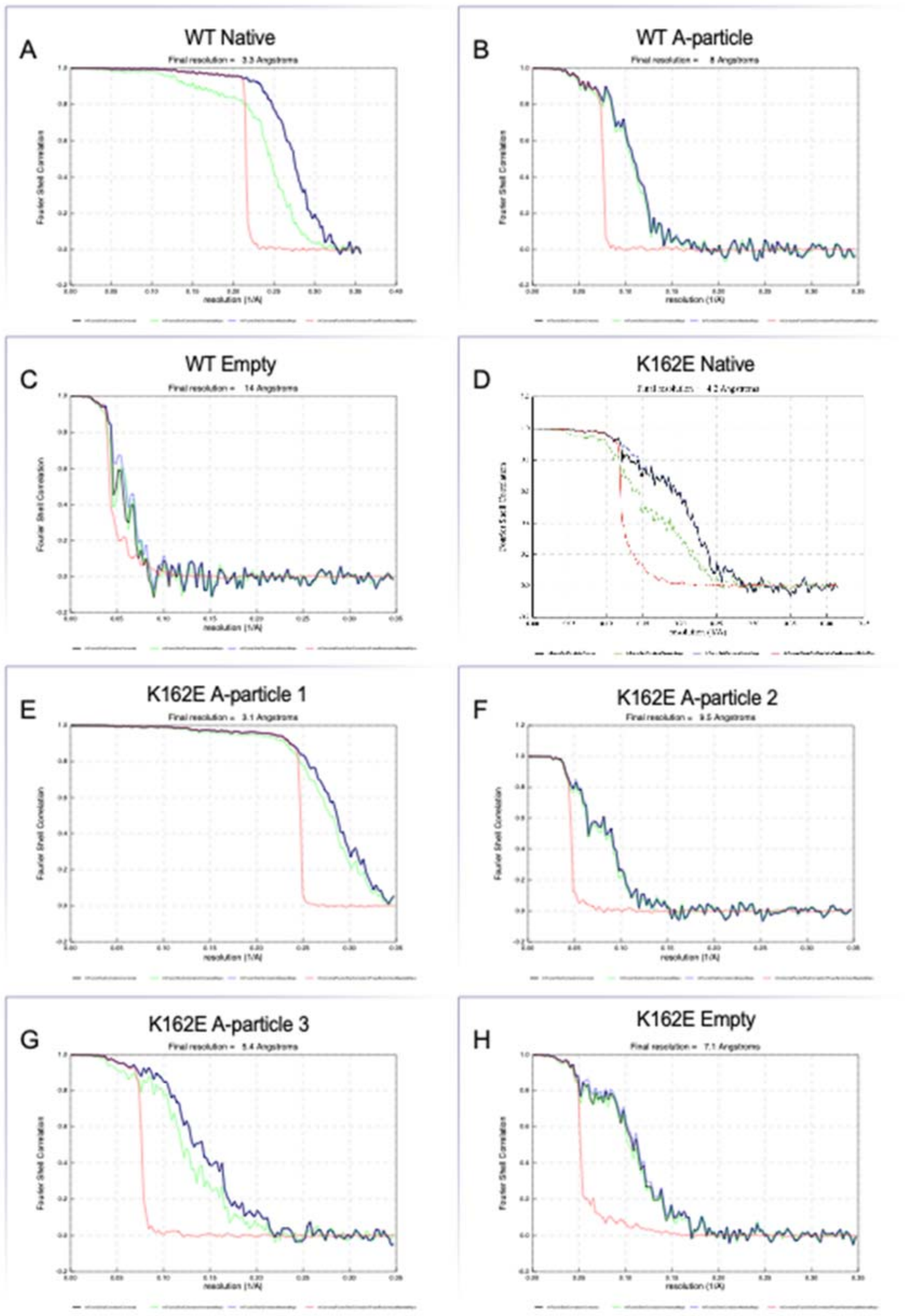


**Fig. 6.** Fourier shell correlations (FSCs) of structures determined. Final resolution calculated by gold-standard 0.143 cutoff of masked maps (blue). Also shown are FSC corrected (black), FSC unmasked (green), and FSC phase-randomized masked (red).

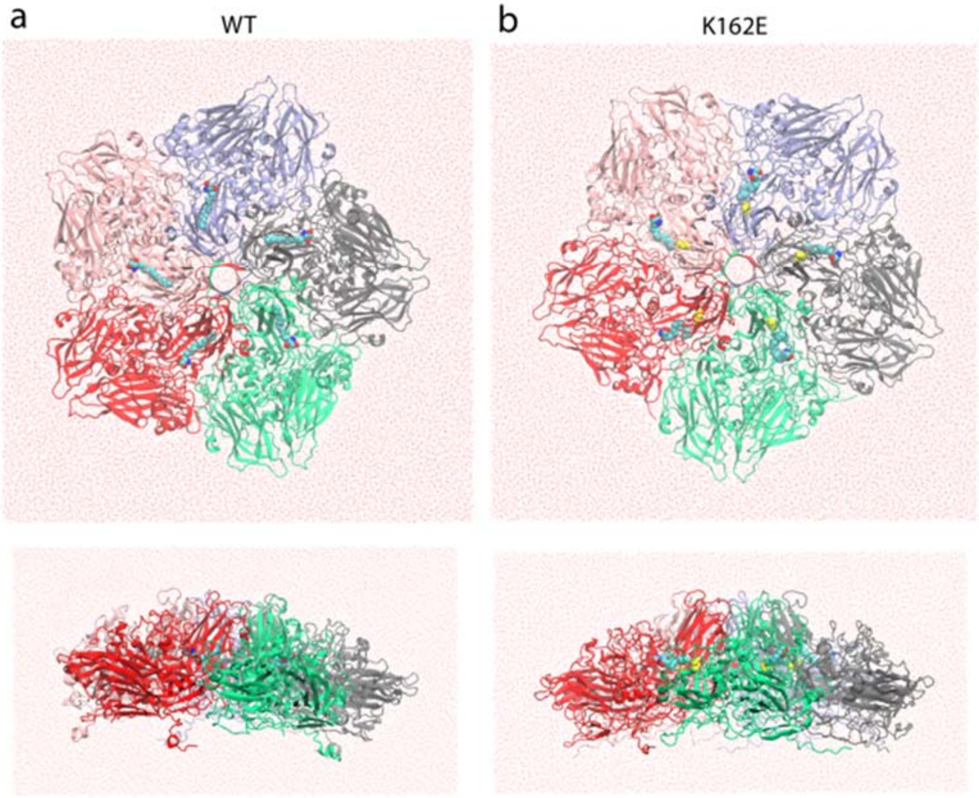


**Fig. 7**. Top and side views of the initial structure of water-simulated box of the WT (a) and K162E (b) capsomeres forming the pentamer. The five capsomeres formed by VP1, VP2, VP3, and VP4 capsid proteins are colored differently. In both systems, sphingosine is represented in van Der Waals format (hydrogen atoms are hidden for clarity) and in the K162E system the VP1 residue 162 is colored yellow and represented in van Der Waals format (hydrogen atoms are hidden for clarity).

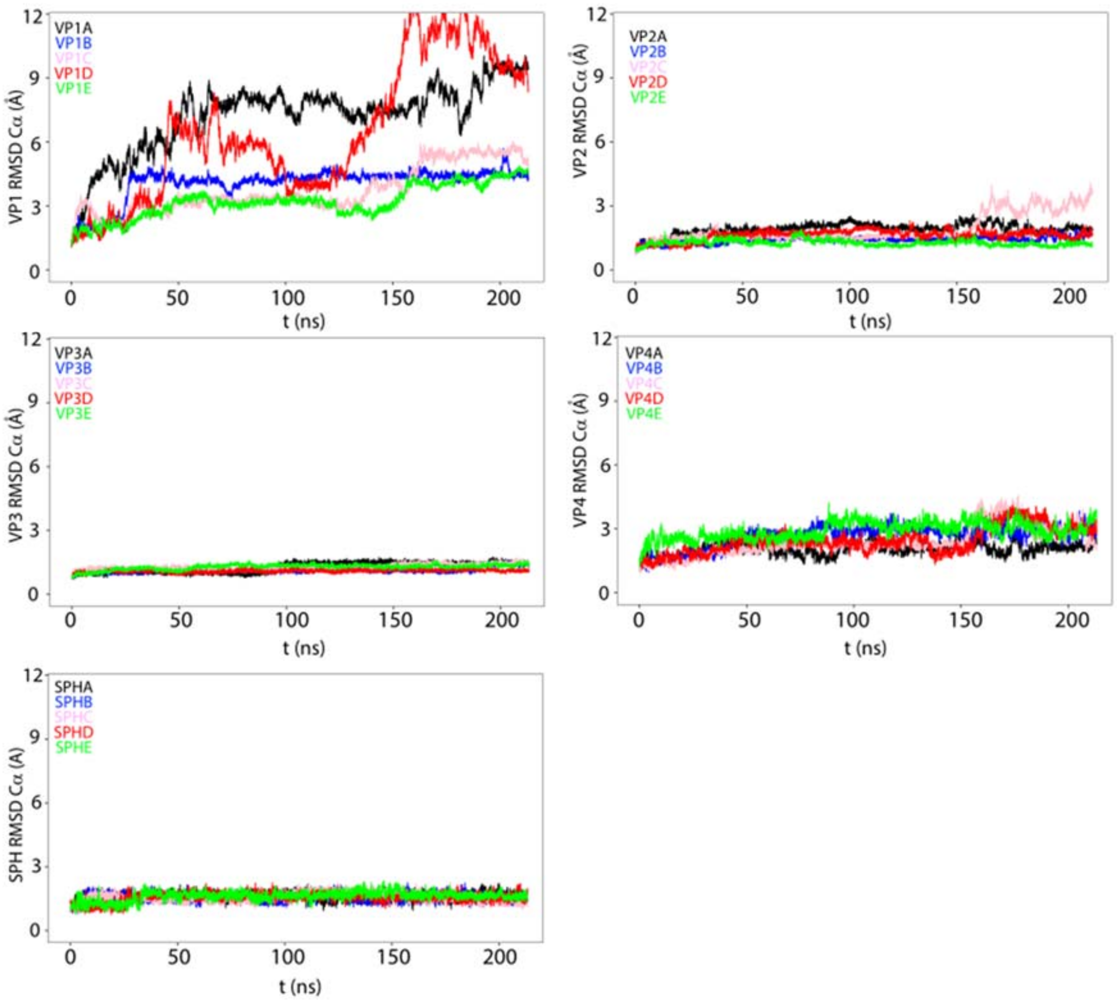


**Fig. 8**. Time evolution of the root mean-squared deviations (RMSD) of the VP1, VP2, VP3, and VP4 capsid proteins and of the sphingosine pocket factor for NPT simulations of the EV-A71 WT carried out at 30ºC. The panels show the time evolution of the RMSD of all the capsid proteins and sphingosine of the five protomers colored differently (protomer A: black; B: blue; C: pink; D: red; E: green).

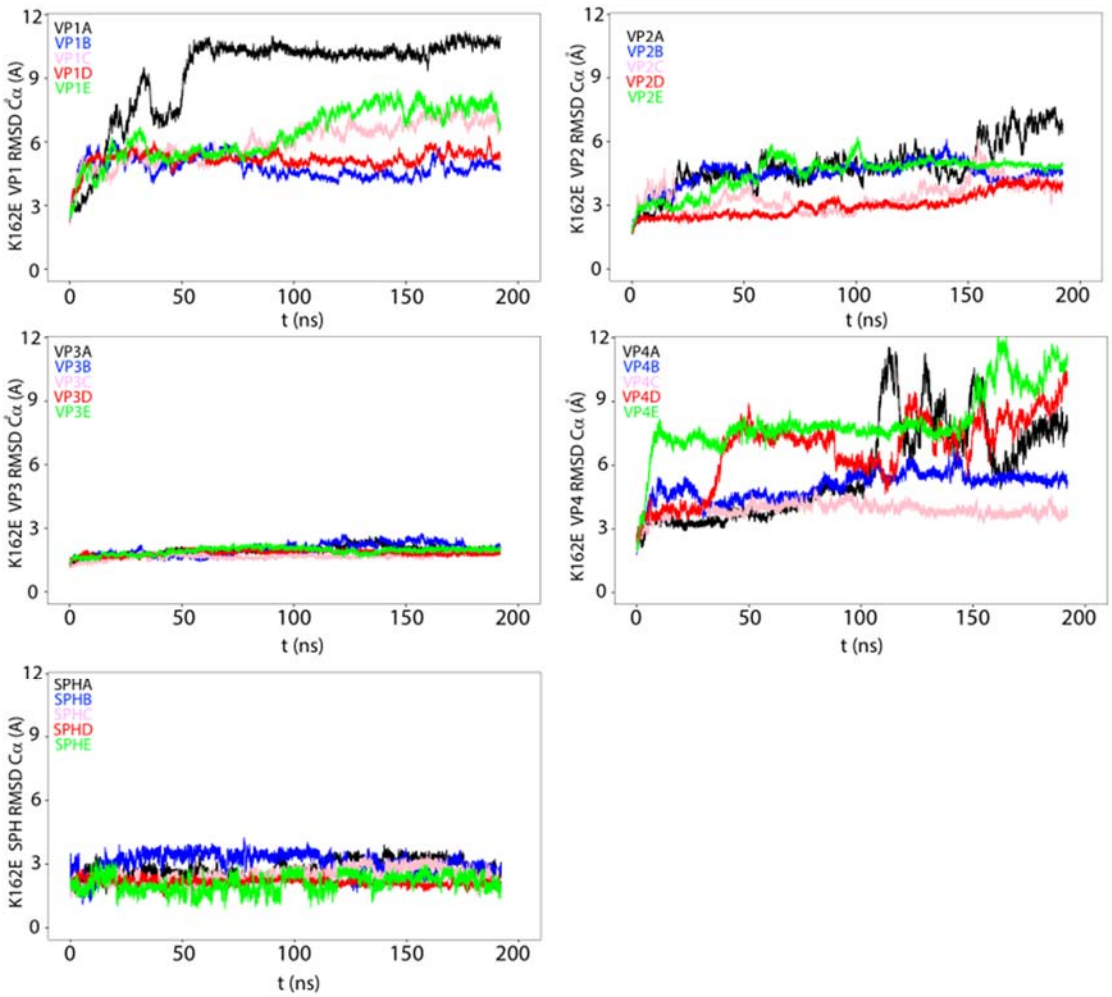


**Fig. 9**. Time evolution of the root mean-squared deviations (RMSD) of the VP1, VP2, VP3, and VP4 capsid proteins and of the sphingosine pocket factor for NPT simulations of the EV-A71 K1162E mutant carried out at 30ºC. The panels show the time evolution of the RMSD of all the capsid proteins and sphingosine of the five protomers colored differently (protomer A: black; B: blue; C: pink; D: red; E: green)

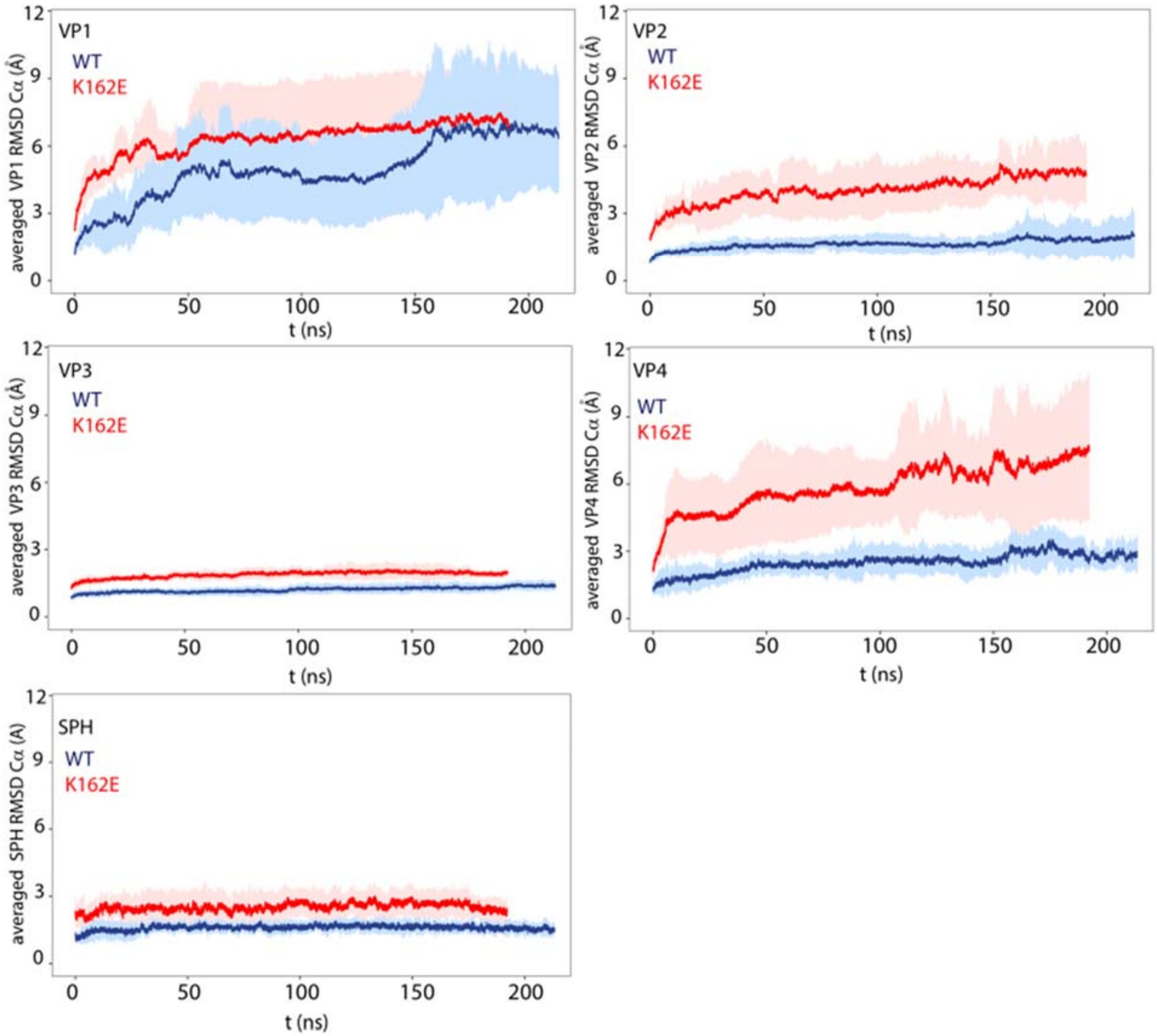


**Fig. 10**. Time evolution of the mean value root mean-squared deviations (RMSD) at 30ºC calculated by averaging the RMSD value at each time point of the simulations of VP1, VP2, VP3, VP4 capsid protein, and of the sphingosine. The error bars represent the standard deviation of the mean.

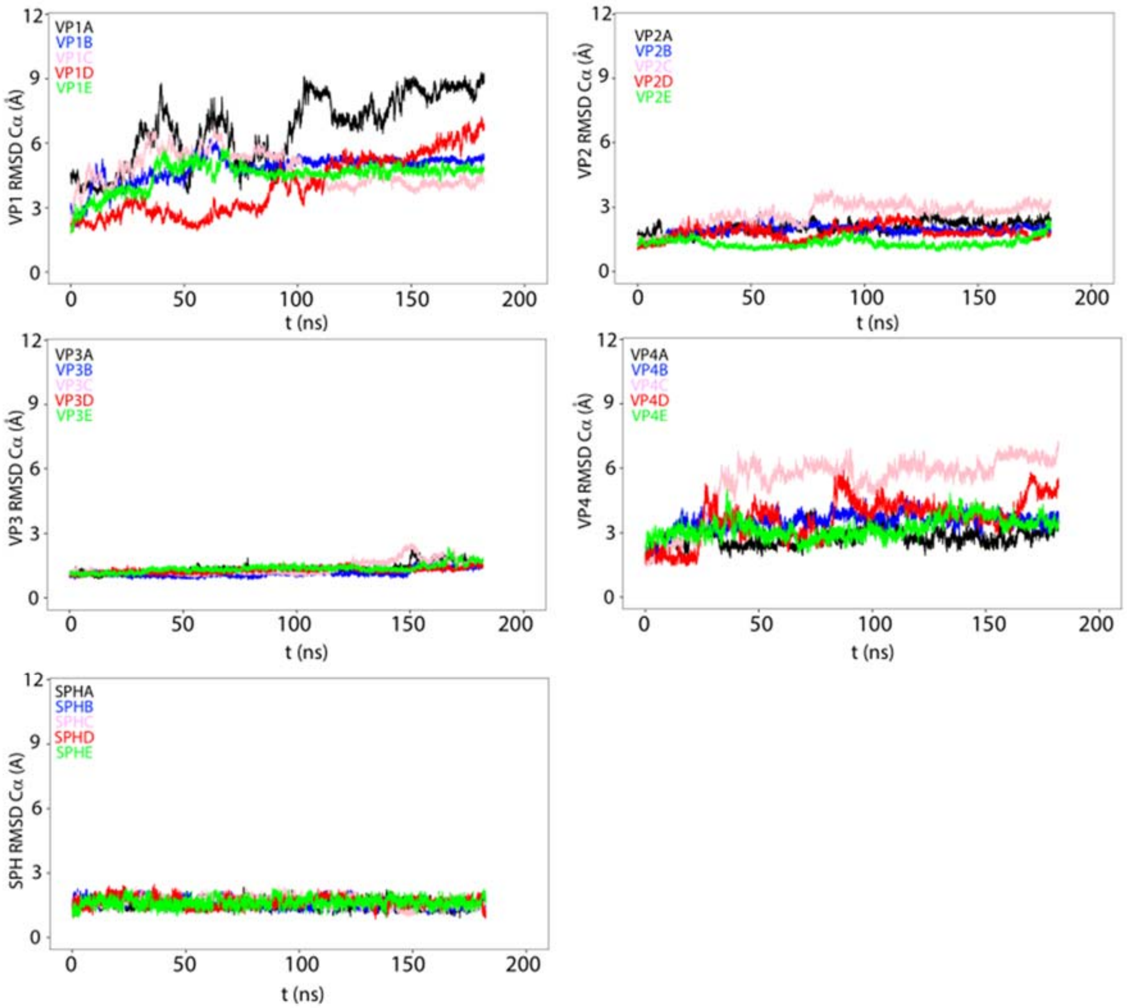


**Fig. 11**. Time evolution of the root mean-squared deviations (RMSD) of the VP1, VP2, VP3, and VP4 capsid proteins and of the sphingosine pocket factor for NPT simulations of the EV-A71 WT carried out at 52ºC. The panels show the time evolution of the RMSD of all the capsid proteins and sphingosine of the five protomers colored differently (protomer A: black; B: blue; C: pink; D: red; E: green)

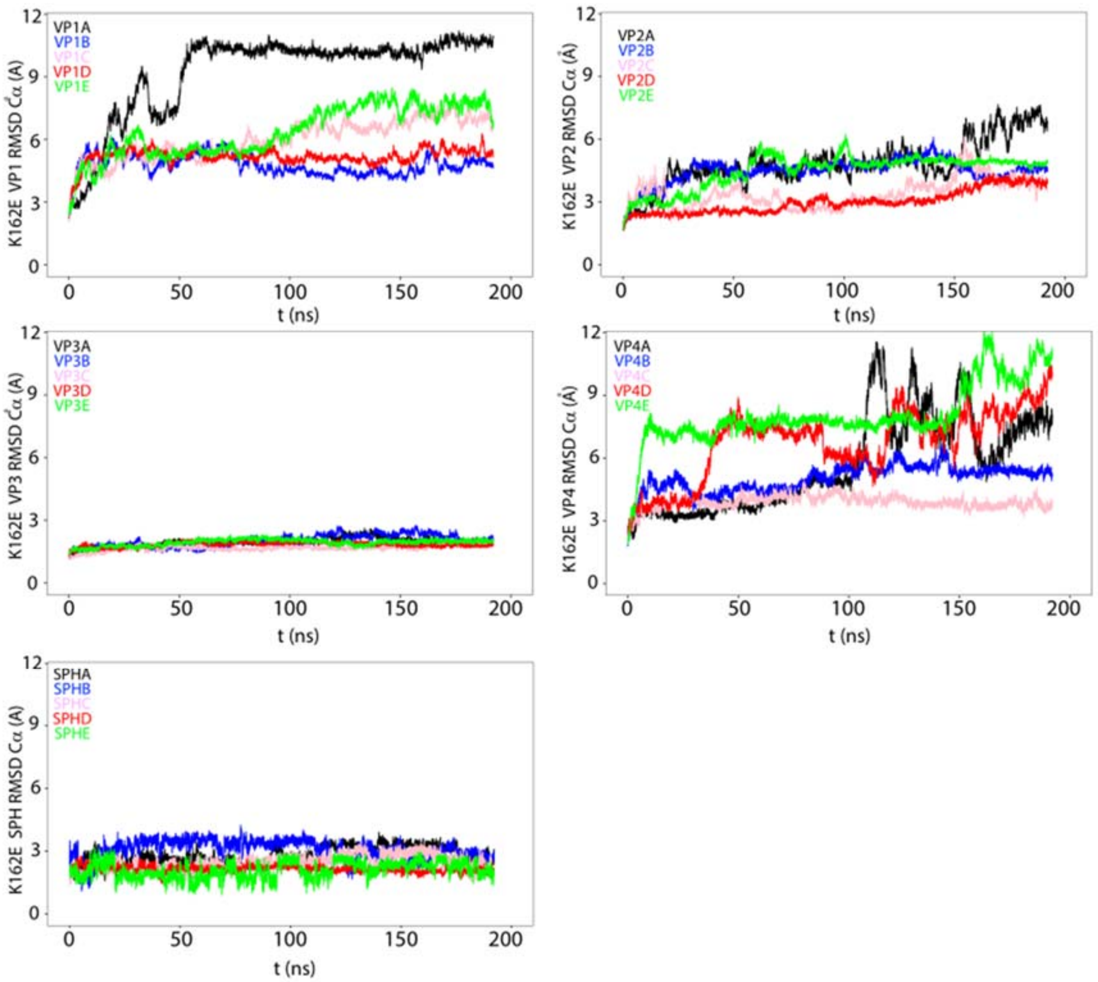


**Fig. 12**. Time evolution of the root mean-squared deviations (RMSD) of the VP1, VP2, VP3, and VP4 capsid proteins and of the sphingosine pocket factor for NPT simulations of the EV-A71 K1162E mutant carried out at 52ºC. The panels show the time evolution of the RMSD of all the capsid proteins and sphingosine of the five protomers colored differently (protomer A: black; B: blue; C: pink; D: red; E: green)

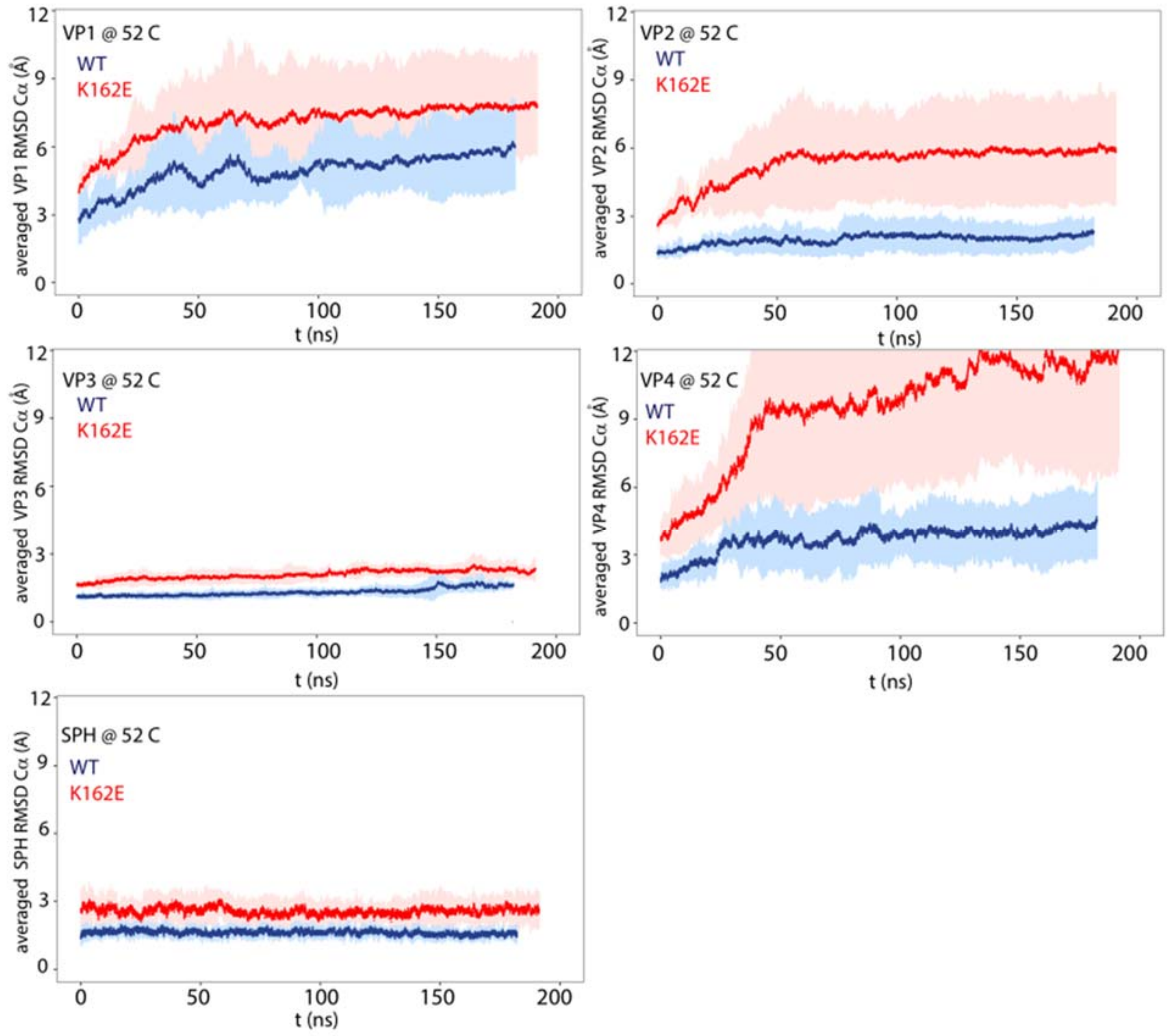


**Fig. 13**. Time evolution of the mean value root mean-squared deviations (RMSD) at 52ºC calculated by averaging the RMSD value at each time point of the simulations of VP1, VP2, VP3, VP4 capsid protein, and of the sphingosine. The error bars represent the standard deviation of the mean.

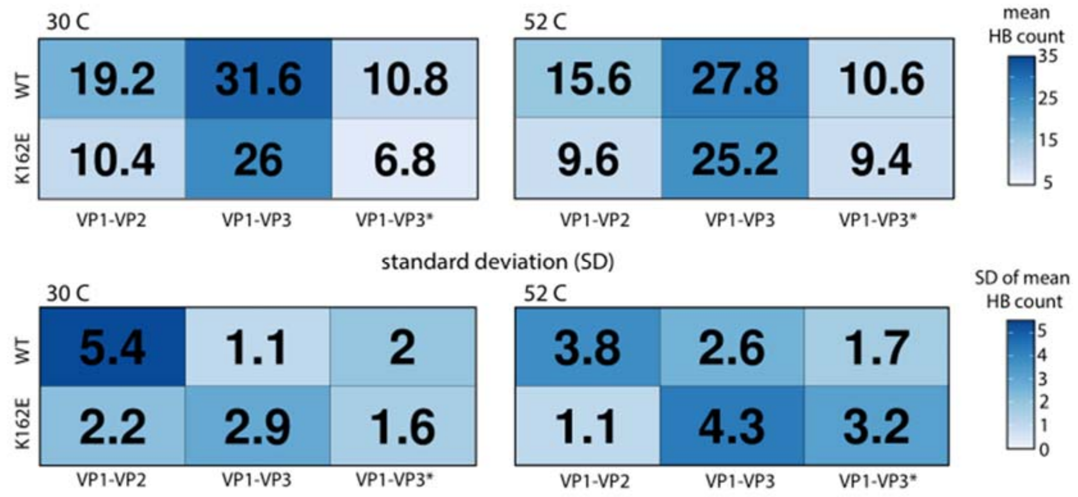


**Fig. 14**. Top panels: Mean of the number of the hydrogen bonds between VP1-VP2 and VP1-VP3 capsid proteins belonging to the same protomer and between VP1-VP3 capsid proteins belonging to adjacent protomers (VP1-VP3*) calculated for the WT and K162E mutant at 30ºC (left panel) and 52ºC (right panel). The calculation has been performed during the last 5 ns of the simulations. Bottom panels: the standard deviation (SD) of the mean of the hydrogen bond (HB) count is represented. The standard deviation is evaluated by considering the five protomers forming the pentamer.

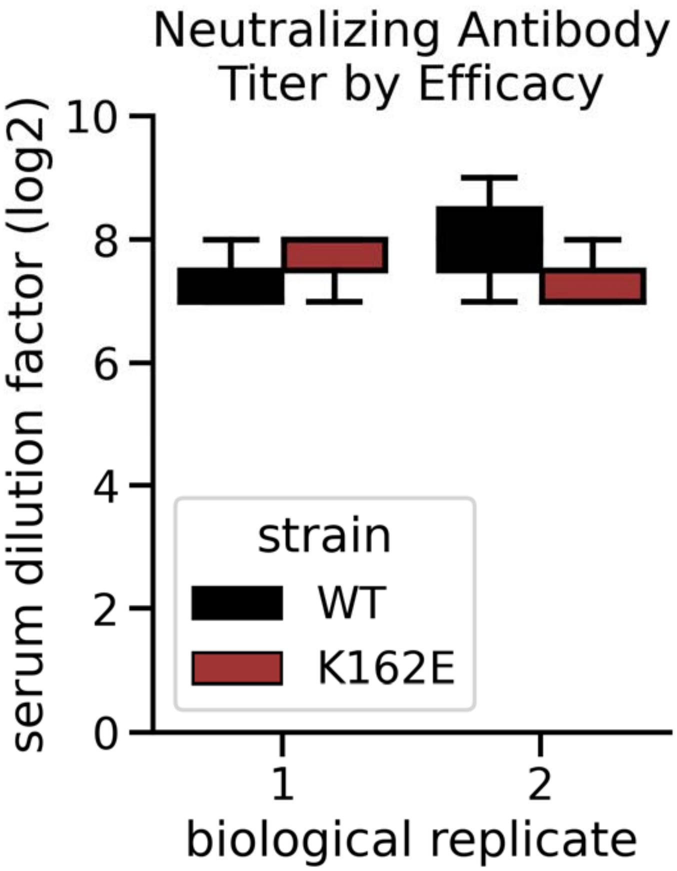


**Fig. 15**. WT and E162K are equally neutralize by polyclonal serum. Either WT or K162E was incubated with different dilutions of serum from a mouse infected with EV-A71. Maximum dilution factor at which neutralization was observed is indicated. Serum was collected from two independent mice and used in biological duplicate, reported dilutions were performed in technical triplicate.

### Movie Legends:

**Supplementary Movie 1. Dynamics of the EF and GH loops in VP1 and VP2 of EV-A71 at 30ºC.** The movie shows the dynamics of the EF loop in green, the GH loop in cyan, and the pocket factor in purple during the length of the simulation compared to their initial position, which is maintained fixed during the length of the movie. For clarity, we maintained fixed also the initial position of VP1, VP2, VP3, and VP4 which are represented in white, new cartoon format.

**Supplementary Movie 2. Dynamics of the EF and GH loops in VP1 and VP2 of EV-**

**A71 K162E thermostable mutant at 30ºC.** The movie shows the dynamics of the EF loop in green, the GH loop in cyan, and the pocket factor in purple during the length of the simulation compared to their initial position, which is maintained fixed during the length of the movie. For clarity, we maintained fixed also the initial position of VP1, VP2, VP3, and VP4 which are represented in white, new cartoon format.

**Supplementary Movie 3. Dynamics of the EF and GH loops in VP1 and VP2 of EV-A71 at 52ºC.** The movie shows the dynamics of the EF loop in green, the GH loop in cyan, and the pocket factor in purple during the length of the simulation compared to their initial position, which is maintained fixed during the length of the movie. For clarity, we maintained fixed also the initial position of VP1, VP2, VP3, and VP4 which are represented in white, new cartoon format.

**Supplementary Movie 4. Dynamics of the EF and GH loops in VP1 and VP2 of EV-A71 K162E thermostable mutant at 52ºC.** The movie shows the dynamics of the EF loop in green, the GH loop in cyan, and the pocket factor in purple during the length of the simulation compared to their initial position, which is maintained fixed during the length of the movie. For clarity, we maintained fixed also the initial position of VP1, VP2, VP3, and VP4 which are represented in white, new cartoon format.
